# Small effect-size mutations cumulatively affect yeast quantitative traits

**DOI:** 10.1101/126409

**Authors:** Bo Hua, Michael Springer

## Abstract

**Summary:** Quantitative traits are influenced by pathways that have traditionally been defined through genes that have a large loss- or gain-of-function effect. However, in theory, a large number of small effect-size genes could cumulative play a substantial role in pathway function, potentially by acting as “modifiers” that tune the levels of large effect size pathway components. To understand the role of these small effect-size genes, we used a quantitative assay to determine the number, strength, and identity of all non-essential genes that affect two galactose-responsive (GAL) traits, in addition to re-analyzing two previously screened quantitative traits. Over a quarter of assayed genes have a detectable effect; approximately two thirds of the quantitative trait variation comes from small effect-size genes. The functions of small effect-size genes are partially overlapping between traits and are enriched in core cellular processes. This implies that genetic variation in one process has the potential to influence behavior or disease in seemingly unconnected processes.

**Highlights:**

- Four yeast quantitative traits are affected by thousands of small effect-size genes.
- Small effect-size genes are enriched in core cellular processes
- The effects of these genes are quantitative trait-specific.

## Introduction

What are all the genes that are involved in a trait? Classically, the pathways that contribute to a trait, like those involved in signaling or development, were defined by genetic screens that identified genes with loss- or gain-of-function phenotypes (Nüsslein-Volhard and Wieschaus, 1980). As screens became more quantitative, many alleles of both small and large effects size where identified (Ehrenreich et al., 2010; Friedman and Perrimon, 2006).But, the methods to validate and then determine the molecular function have remained laborious. Hence, research has typically focused on genes on characterizing genes with large effect size. This has lead to a potential bias that these large effect size genes dominate the behavior and variability in a pathway. An alternative view is that cumulatively, the mainly overlooked small effect size genes significantly shape pathway function and population-level trait variation, and hence the genetic architecture of a pathway is distributed not centralized (Figure 1). Until recently, it wasn’t possible to easily and comprehensively identify genes implicated in quantitative traits, making it difficult to distinguish between these two hypotheses concerning the architecture of most pathways.

**Figure 1.**
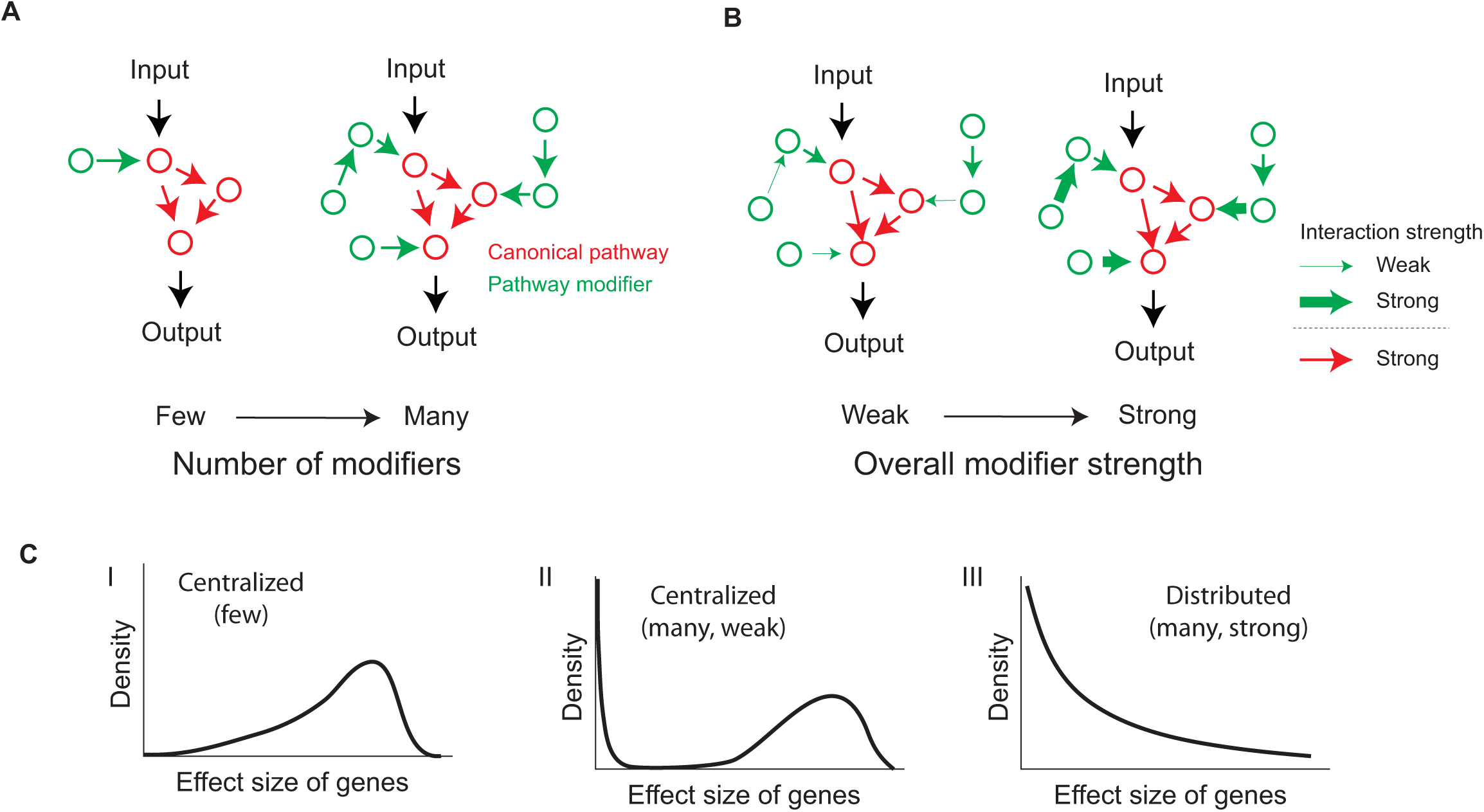
Genes outside of the canonical signaling pathways have the potential to substantially influence pathway function. (**A**) A canonical pathway (red circles) can be modified by anywhere from a small number to large number of currently unidentified genes (green circles). (**B**) Regardless of the number of modifiers, the modifiers could range from having a weak to strong effect on the pathway (represented by arrow thickness). (**C**) If the number of modifiers is small (I) or if the effect size of the modifiers is small (II) the genetic architecture of the pathway will be centralized, i.e. a small number of genes will control the function of and variation in the pathway. If the number of modifiers is large and the effect size of the modifiers is sufficiently large (III) the genetic architecture will be distributed; i.e. a large number of genes will control the function of and variation in the pathway.

The genetic architecture of quantitative traits has taken on increased importance as it has become clear that many human traits, such as body mass index and traits that underlie heritable human disease, are also quantitative. Numerous human traits and disease have been studied using genome-wide association studies (GWAS) to uncover the loci containing causative variants that are responsible for the genetic component of these traits (Hindorff et al., 2009). If the genetic architecture of the underlying pathway were centralized, one would expect GWAS would yield a small number of large effect-size genes typically of related function; if the genetic architecture of a quantitative trait were distributed, one would expect GWAS would yield a large number of small effect-size genes of often seemingly unrelated function. In some diseases, e.g. age-related macular degeneration (AMD), GWAS indeed identified several common alleles of large effect size that explain about half of the disease risk to siblings of affected individuals (Maller et al., 2006). This would support the view of centralized signaling pathways. But, in many cases, GWAS has yielded many small effect-size variants with low odd ratios (Hindorff et al., 2009), and additionally many identified loci have not included genes with an obvious connection to disease (Cooper and Shendure, 2011; Edwards et al., 2013). These results are consistent with the hypothesis that the gene architecture of some pathways underlying human traits is distributed. Direct experiments to separate between these two hypotheses can help frame our expectation for the results from these association studies.

Model organisms should be a powerful set of tools for defining the architecture of quantitative traits. Several studies in yeast (Bloom et al., 2013; Ehrenreich et al., 2010) show that linkage analysis has the potential to identify most of the causative loci needed to explain trait variation between two natural yeast isolates. But these studies, like human GWAS, are limited by recombination block size and sample size, and hence are not ideal for identifying causative genes or the exact number and identity of small effect-size loci. As an alternative approach, deletion libraries have been used to assess the role of every yeast gene. These studies have been transformative for defining the function of unknown genes (Botstein and Fink, 2011) and for showing that many processes in yeast are genetically interconnected (Costanzo et al., 2010; 2016). While informative, the assays that are typically performed, e.g. colony size assay, are not quantitative enough to accurately determine the effect size of every mutant. Hence whether this interconnectedness has a significant role in pathway function is still unclear.

In this work, we quantified the effect sizes of all non-essential yeast genes on several traits. Instead of identifying existing genetic variation in natural populations, we used a yeast deletion library to measure with high precision the magnitude of effect of all non-essential genes on a quantitative trait, which we refer to as gene effect size. By its design, this approach identifies all the genes whose loss-of-function has the potential to influence a trait, and the effect size distribution of these genes. We found that all four traits we analyzed have an exponential distribution of effect sizes. The consequence of these results is that cumulatively, small effect-size can significantly contribute to pathway function. Gene Ontology(GO) analysis and additional experiments showed that many of these small effect-size mutations are involved in core cellular processes and affect quantitative traits in a trait-specific, not generic, manner. In natural populations, phenotypic variation is influenced by the actual existing variants; this natural variation is more complex than our deletion library. We showed through simulation that our analysis based on deletion mutants, given modest assumptions, yields an effect size distribution that is close to the distribution that would be observed for other sources of genetic variation.

## Results

### A large fraction of genes can influence multiple biological processes

A large number of screens have been performed with the yeast deletion library (Giaever and Nislow, 2014). These screens could potentially serve as a rich source of data for determining the effect size of each gene on many traits. Reanalyzing this data, we found that, due to measurement noise, most of these studies do not have the power to determine the full gene-level effect size distribution (Supplemental Information). This is not surprising as the goal of most studies was to identify genes of large effect size rather than attempting to identify all genes of any effect size. Therefore, to determine the number of genes that can affect a pathway, we created a reporter library with which we could quantitatively measure the response of cells to galactose (GAL). We systematically constructed a library of strains deleted for all non-essential yeast genes each containing a YFP reporter driven by the GAL1 promoter (GAL1pr-YFP). We then assayed the bimodal YFP response (Acar et al., 2005; Escalante-Chong et al., 2015) in single cells growing in mixtures of glucose and galactose by flow cytometry (Figure 2A-B). Additionally, to supplement the analysis, we identified and re-analyzed two deletion studies (Breslow et al., 2008; Jonikas et al., 2009), one on growth rate in rich medium and one on the unfolded protein response (UPR), that had a signal-to-noise ratio that was sufficiently large to determine the effect size distribution.

**Figure 2.**
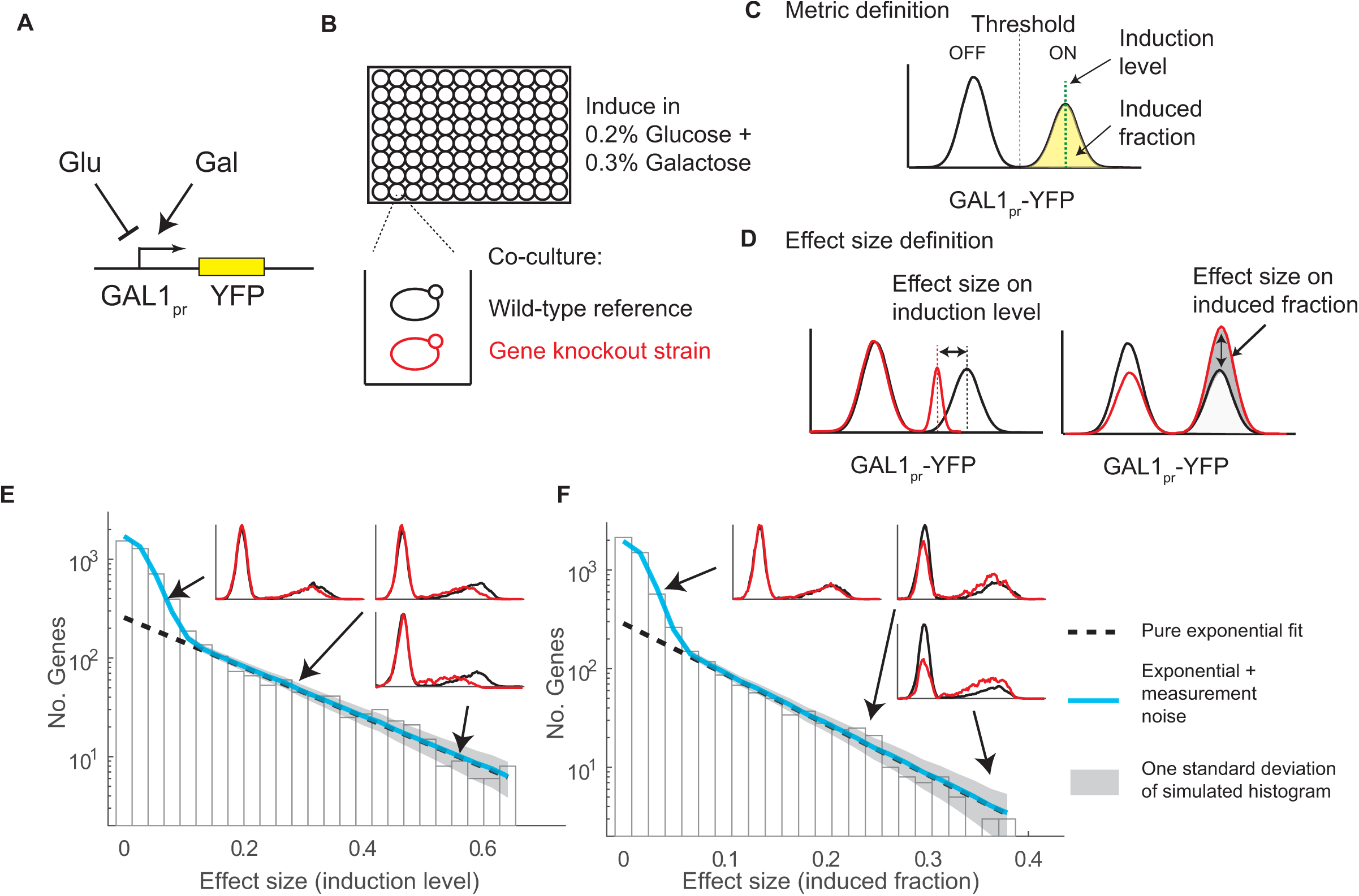
Quantitative genetic screen determines that a large number of genes quantitatively affects the yeast galactose response. (**A**) Galactose (Gal) activates while glucose (Glu) inhibits transcription from a GAL1 promoter YFP fusion. **B**) A mCherry expressing mutant strain (red) was co-culture with a wild-type reference strain (black); both strain contained the reporter construct from **A**. Each well contained a distinct deletion mutant. (**C**) We defined two metrics to characterize the bimodal response of the GAL pathway. We defined the induced fraction (yellow area versus total area under the curve) as the percent of cells whose YFP expression level was above a threshold (black dotted line). We also defined the induction level as the mean YFP expression of all induced cells (green dotted line). (**D**) Mutant effect sizes for the induction level (**D**, left) and for induced fraction (**D**, right) are defined as the relative change in each metric between mutant (red) and the co-cultured wild-type reference strain (black). (**E**-**F**) Effect size distribution for two GAL traits. Effect sizes of all mutants were binned and plotted as a histogram (black bars). Mutant that passed a 0.5% false discovery rate cut-off were well fit with an exponential distribution using maximum likelihood estimation (dashed black line, R^2^ 0.96 for each, see Method). The full distribution is parsimonious with a convolution of experimental noise and an exponential distribution (blue line is the average distribution of 100,000 simulations, R^2^=0.92-0.98; gray shading is one standard deviation around the mean).

Principal component analysis of the results from our GAL response screen highlighted three distinct traits (Figure S1). These traits, corresponding to: 1) the fraction of cells that are induced above background; 2) the induction level of the induced ('on') peak; and 3) the background level of the uninduced ('off') peak (Figure 2C, Supplemental Information). The signal-to-noise ratio of the first two metrics was sufficient to calculate an effect size distribution for a large number of genes. We will refer to these two separable GAL traits as the "induced fraction" and the "induction level" (Figure 2D).

Each of the four traits - the induced fraction, induction level, growth rate, and UPR - considered in isolation, was influenced by a large number of deletion strains (Figure 2E and F); the distribution of mutant effects was continuous. Based on a comparison of the measured effect sizes and the measurement noise estimated from biological replicates, 19% (796 of 4201), 16% (735 of 4562), 16% (689 of 4162), and 20% (849 of 4162) of non-essential genes screened, at a 0.5% false discovery rate, affect the growth rate in rich media, unfolded protein response, induced fraction in GAL, and induction level in GAL respectively. Together the two GAL traits are composed of 1104 unique genes. Interestingly, if we used a single composite trait, i.e. mean expression, to quantify the GAL response, fewer genes (593 of 4162) were identified, highlighting the utility of sub-classifying higher-level phenotypes that might be composed of separable traits each controlled by distinct genetic factors (Supplemental Information). To obtain a more accurate estimate of how many genes can quantitatively affect each of the traits, at the sacrifice of knowing the identity of the genes, we determined the area of the normalized effect size distribution that is outside the normalized measurement noise distribution (Figure S2). From this, we estimate that the fraction of genes affecting the growth rate in rich media is 62%, unfolded protein response is 23%, induced fraction in GAL is 28%, and induction level in GAL is 34% (Supplemental Information). Together these results highlight that a large fraction of the protein-coding genes has the potential to quantitatively affect a trait.

As a final method to determine the number of genes that influence our four traits we determined whether the effect size distributions could be explained by a simply analytical function. To minimize the effect of measurement noise on measured effect sizes, we first focused our analysis on the genes whose effect size was significantly different from measurement noise. Interestingly, we found that the effect size distribution for all four traits was well fit by an exponential distribution (R^2^=0.91-0.96, Figure 2E an F, dotted line). When extrapolating the exponential fit into the measurement noise, it predicts that 27-33% of genes affect each of our four traits, similar to the orthogonal estimates above. Adding measurement noise to the exponential distribution (Figure 2E and F, blue line) well fit the full measurement distribution (R^2^=0.92-0.98). Therefore, a parsimonious explanation of our data is that the effect size distribution of a quarter to half of genes is exponential. Half to three quarters of all genes have little to no effect.

### Small effect-size genes can influence pathway function

The shape of the determined effect size distributions implies that each of the four traits is affected by genes with a continuous distribution of effect sizes ranging from a small number of large effect-size genes to a large number of small effect-size genes. It has been questioned whether even such a large number of small effect-size mutants could substantially contribute to the functionality of a pathway (Crow, 2011). The answer to this question depends on the exact shape of the measured effect size distribution (e.g. Figure 1C II versus III). We therefore determined the number of genes that are cumulatively important for pathway function. To do so, we devised a method to quantify the impact of each gene, which is similar to the one used to quantify allelic contribution to narrow-sense heritability in a GWAS (Lynch and Walsh, 1998). In the calculation, we first assumed a population of cells with independent and randomly assorting alleles. We assumed only two possible alleles for each gene, i.e. deletion or wild-type (a more complex model will be considered below). We then calculated each gene’s contribution to the trait variation in the population as 2*β*^2^*f*(1 – *f*) (Lynch and Walsh, 1998), where *β* is the effect size and *f* is the allele frequency, assuming that each allele has a frequency of 50% and no epistasis (Figure 3A). For our four traits, using the measured effect size for each gene, we find that 257-352 genes with the largest effect sizes, representing 5.6-8.5% of screened genes, are needed to explain 80% of total computed variation (Figure 3B and C, and Figure S4). If human traits behave similarly to our yeast deletions, we would estimate that the number of genes required to explain most of the heritability of a quantitative trait is in the range of 1200-1900 genes. Interestingly, our estimate is concordant with estimations from GWAS. For example, the current estimate for human height, the best characterized human trait, is that 423 1Mb loci are involved. Yet this explains only 20% of the heritability. This result suggests that both in yeast and humans, some pathways and traits resemble the distributed architecture from Figure 1C III; i.e. a large number of genes of slowly diminishing effect size contribute to pathway function and trait variation.

**Figure 3.**
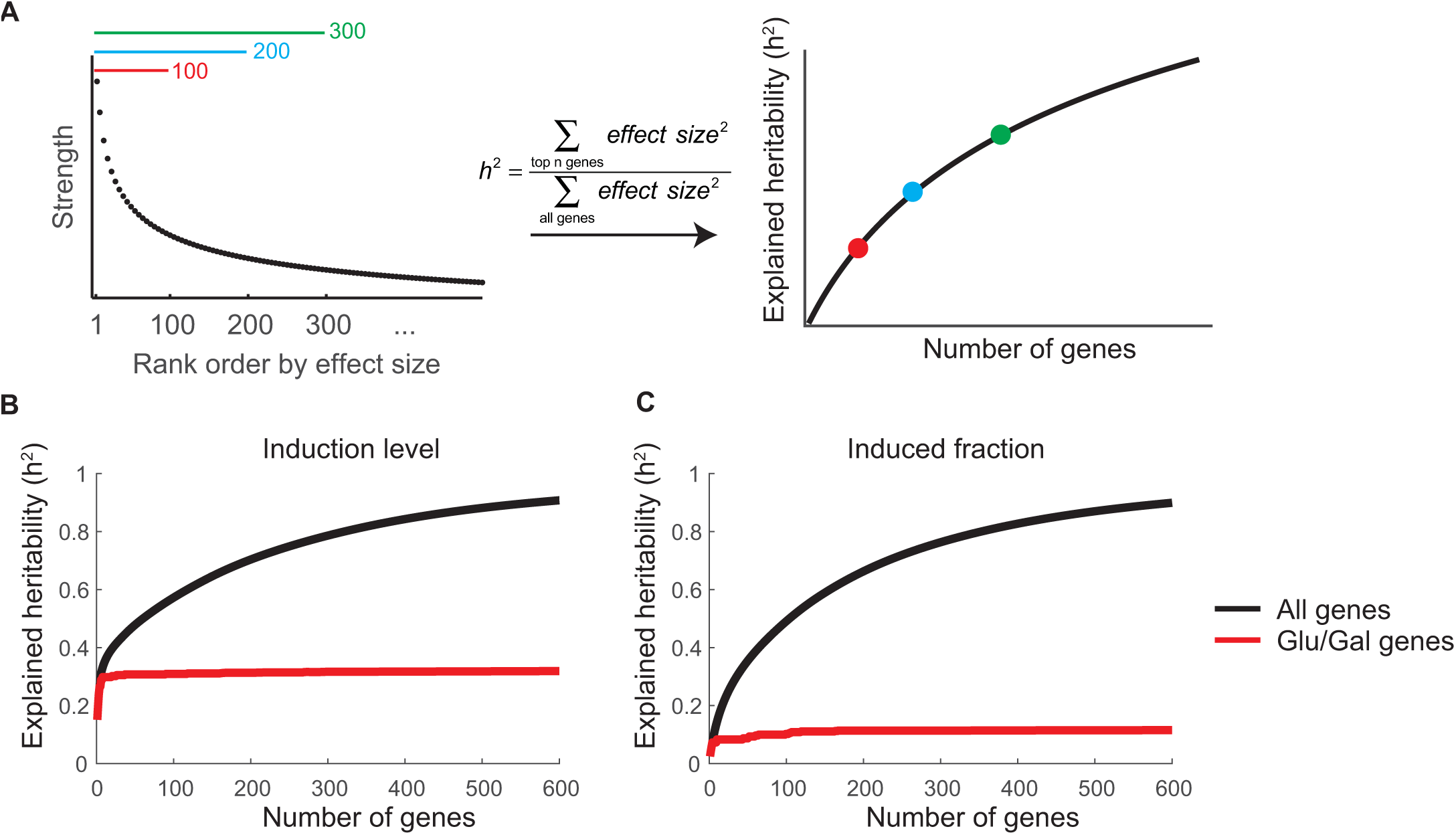
Pathway modifiers can significantly contribute to heritability. (**A**) Methods to estimate the heritability explained by a set of deletion mutants. Genes were sorted based on their effect size when deleted. The heritability was calculated as the sum of the squares of the effect sizes for the top n genes compared to all genes. The heritability (right) for the top 100 (red), 200 (blue), and 300 (green) mutant strains (left) is shown. (**B-C**) The contribution to explained heritability, as calculated in **A**, from GAL genes (red) or all genes (black) for induction level (**B**) and induced fraction (**C**).

Given their individual small effect size, our analysis also suggests that a significant portion of the genes that account for pathway function would not typically be considered to be contributing to each trait. Classical genetic screens identified only a fraction of the genes that have the potential to significantly affect each of the two GAL traits. A compiled list of the 50 genes previously identified as affecting the GAL pathway (Supplemental Information) explained only 32.0% and 11.7% of variation in the induction level and induced fraction traits respectively. Similarly, in the unfolded protein response, genes whose products localize throughout the secretory pathway (ER and Golgi) explain only 27.1% of variation, further suggesting substantial roles of additional genes/processes. Hence, much of the variance occurs in genes we term non-trait-specific, i.e. genes that are not typically considered to be physiologically related to the trait. This is consistent with previous GWAS that identified putative causative loci that in some cases contained genes that were obviously trait-specific but in other cases were involved in general cellular processes. For example, human height is affected by variants in genes that underlie skeletal growth defects (trait-specific), as well as general pathways such as the Hedgehog pathway (non-trait-specific) (Lango Allen et al., 2010). Surprisingly, our analysis suggests that the non-trait-specific processes can have a larger aggregate effect than trait-specific pathways.

A potential caveat to these estimates is that the GAL phenotypes of ten mutants are either fully induced or uninduced, causing the effects of these genes to be underestimated. These ten genes have previously described influences on the GAL pathway. *GAL1, GAL3, GAL4, GAL80, REG1*, and *SNF3* are involved in either glucose or galactose signaling. *HSC82* and *STI1* interact with the *HSP90* co-chaperone that has been shown to influence the GAL pathway (Gopinath 2016). *SNF2* is a SWI/SNF chromatin remodeling complex that was previously suggested to be involved in nucleosome occupancy on GAL promoter (Bryant et al., 2008). *GCN4*, is a general transcription factor that responds to amino acid starvation. We believe in most cases this caveat does not affect our results. Because, the loss-of-function effect size of these alleles is effectively infinite, they will behave as Mendelian not quantitative alleles. Instead, for any quantitative traits, the predominant alleles of these Mendelian loss-of-function genes must be hypomorphic alleles. Indeed, when we assume hypomorphic allele effect sizes for these genes by randomly sampling from the tail of the fitted exponential distribution, we only observed a modest increase in total trait variation (< 3%).

### Gene deletions in core cellular processes affect quantitative traits

What are the functions of these ‘pathway modifiers’ we identified? Are they genes that have general effects on all traits or are they specific to one or a subset of traits? We found that non-trait-specific processes often affect more than one trait. All pairs of traits share significantly more genes that affect their behaviors than expected (p < 10^-65^, one-tailed hypergeometric test). While only 2 genes would be expected by chance, 113 genes were shared by all traits (Figure 4A). These genes also overlap significantly with "hub" genes identified from genetic interaction network (between 140 and 257 out of 380 hubs genes are significant for each of the four traits, p < 10^-47^, hypergeometric test) (Costanzo et al., 2010). We used Gene Ontology to ask if shared non-trait-specific genes were enriched for specific biological processes. Indeed, many processes were enriched (Table S1), including translation (GO:0006412), regulation of metabolism (GO:0031323), and transcription (GO:0006351). Although this has not previously been extensively characterized, it is not surprising that these traits might be altered by perturbations in some core cellular processes.

**Figure 4.**
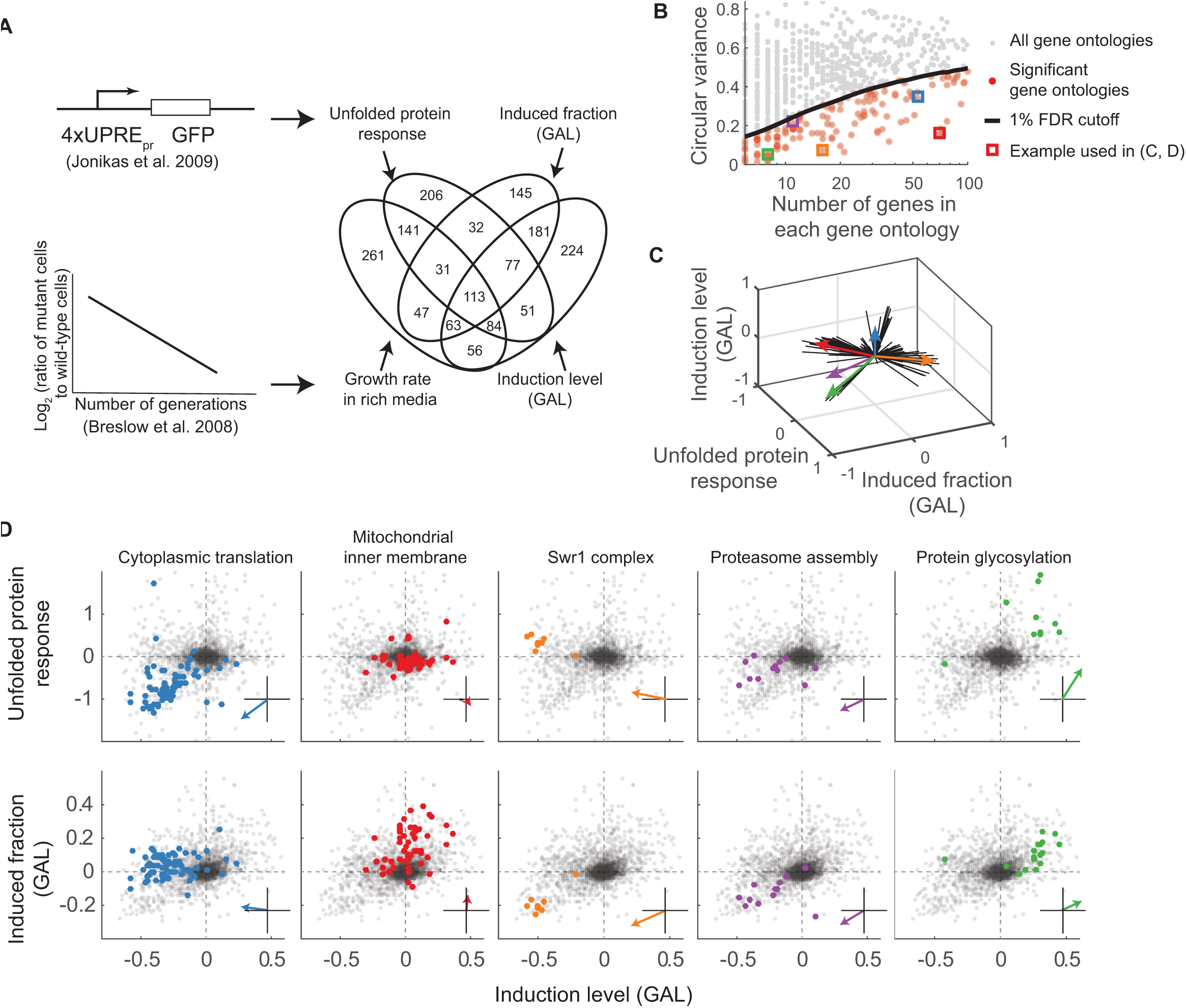
Core cellular processes affect quantitative traits. (**A**) Venn diagram showing the overlap between genes that significantly affect each of our four quantitative traits. Effect size for the unfolded protein response and growth rate in rich media was determined by reanalyzing data from Jonikas et al. and Breslow et al. (Breslow et al., 2008; Jonikas et al., 2009). Only genes that were assayed for all four traits are included in the Venn diagram. (**B**) Identification of gene ontologies (GOs) that are significantly clustered in the 4-D trait space. For each GO the mean circular variance in the 4-D trait space was determined (Methods) and plotted against the corresponding number of genes in that GO (orange dots are significant, gray dots are not). To determine the 1% false discovery rate (FDR < 1%, black line), gene names were permuted (10000 bootstraps) before calculating the circular variance. GOs displayed in **C** and **D** are shown as squares. (**C**) The average effect size vector for each significant GO in **B** projected into the 3-D induced fraction-induction level-UPR response space. (**D**) Examples of GO with distinct spatial clustering. The effect of gene deletion on the unfolded protein response vs. GAL induction level (top) and on the GAL induced fraction vs. GAL induction level (bottom) was plotted for all genes from five different significant GOs from **B** (GO genes in color, all other genes in gray). (Inset) Average mutant vector of GO.

The identification of these core cellular processes as having potential to explain a significant amount of trait variation could be fundamental or trivial. It could reflect an architecture where many biological traits integrate many external and internal factors as inputs (e.g. the GAL pathway responding not just to galactose but glucose, redox status, ribosome capacity, ER capacity, etc.). Alternatively, as the expression of a large fraction of yeast genes is affected by growth rate control (Keren et al., 2013; Regenberg et al., 2006; Slavov and Botstein, 2011), a trivial explanation could be that the effect on the UPR and GAL traits is solely an indirect effect of a growth rate defect (Figure S4). Our data do not support growth rate as the sole factor explaining our results. Between 40% and 60% of gene deletions affect our GAL and UPR traits without affecting growth and vice versa (Table S2). Furthermore, for genes that affect both growth rate and any of the other traits, there is no correlation in effect size between the two effects (R^2^<0.02, S Figure 7B-D). These observations argue against the idea that defects in growth are the main reason that non-trait-specific genes affect the behavior of traits. The involvement of many non-trait-specific genes instead suggests that many signaling pathways integrate a much larger set of cellular inputs than the single input for which the pathways are named.

### Perturbation of core cellular processes can have trait-specific effects

Consistent with the idea that traits integrate a number of inputs in a trait-specific manner, we find that biological processes often affect more than one trait, but importantly not all traits. Using a spatial clustering algorithm in the four-trait space (Figure 4B-C), we found an enrichment in core cellular components (Table S3), such as ribosomal genes (GO:0002181), mitochondrial genes (GO:0005743), mannosyltransferases (GO:0000030), genes that affect histone exchange (GO:0000812) or proteasome assembly (GO:0043248), and genes involved in peptidyl-diphthamide synthesis (GO:0017183). Each of these sets of genes had a separable direction in this 4-dimensional space suggesting each process is responding distinctly to the mutations (Figure 4D). For example, the 89 genes involved in cytoplasmic translation (GO:0002181) were enriched in 3 out of 4 quantitative traits, namely the unfolded protein response, growth rate in rich media, and GAL induction level, but not GAL induced fraction (40, 21, 37 and 3 genes respectively out of the top 300 genes). Conversely, mitochondrial inner membrane genes (GO:0005743) are enriched in the GAL induced fraction but not growth rate, unfolded protein response, nor GAL induction level (21 versus 3, 1, and 4 respectively out of the top 300 genes).

Furthermore, the same core processes can have distinct effects on different traits. For example, at first glance, one might expect mutations in ribosomal genes to affect the level of induction of a pathway (e.g., by altering the expression level of all genes) but not the fraction of cells induced. Indeed, this is the case for the GAL response. But, when we examined the effect of the same mutants on a phosphate responsive (PHO) promoter, PHO84pr, we obtained a different result (Figure 5). The *PHO84* promoter responds to phosphate limitation in a bimodal manner and can therefore be characterized in the same way as we characterize the GAL response. The effects of ribosomal mutants on the induction level of PHO84pr-YFP are significantly less than for GAL1pr-YFP (Figure 5B versus C; Figure 5E and Figure S5, p=3x10^-10^, two-tailed t-test). Instead, ribosomal mutants affect the PHO induced fraction and the level of expression of the uninduced cells (Figure 5C and examples in Figure 5D). In support that these results are a direct consequence of perturbation of ribosomal function, cycloheximide, a small molecule inhibitor of the ribosome, phenocopies the results of ribosomal gene deletions on both the GAL and PHO pathways (Figure 5B-D). While this result at first may seem counter-intuitive, these results could be explained if ribosomal proteins differentially impacted the expression level of positive versus negative regulators of a trait. In total, this suggests that variation in genes involved in core cellular processes could have both generic and pathway specific effects.

**Figure 5.**
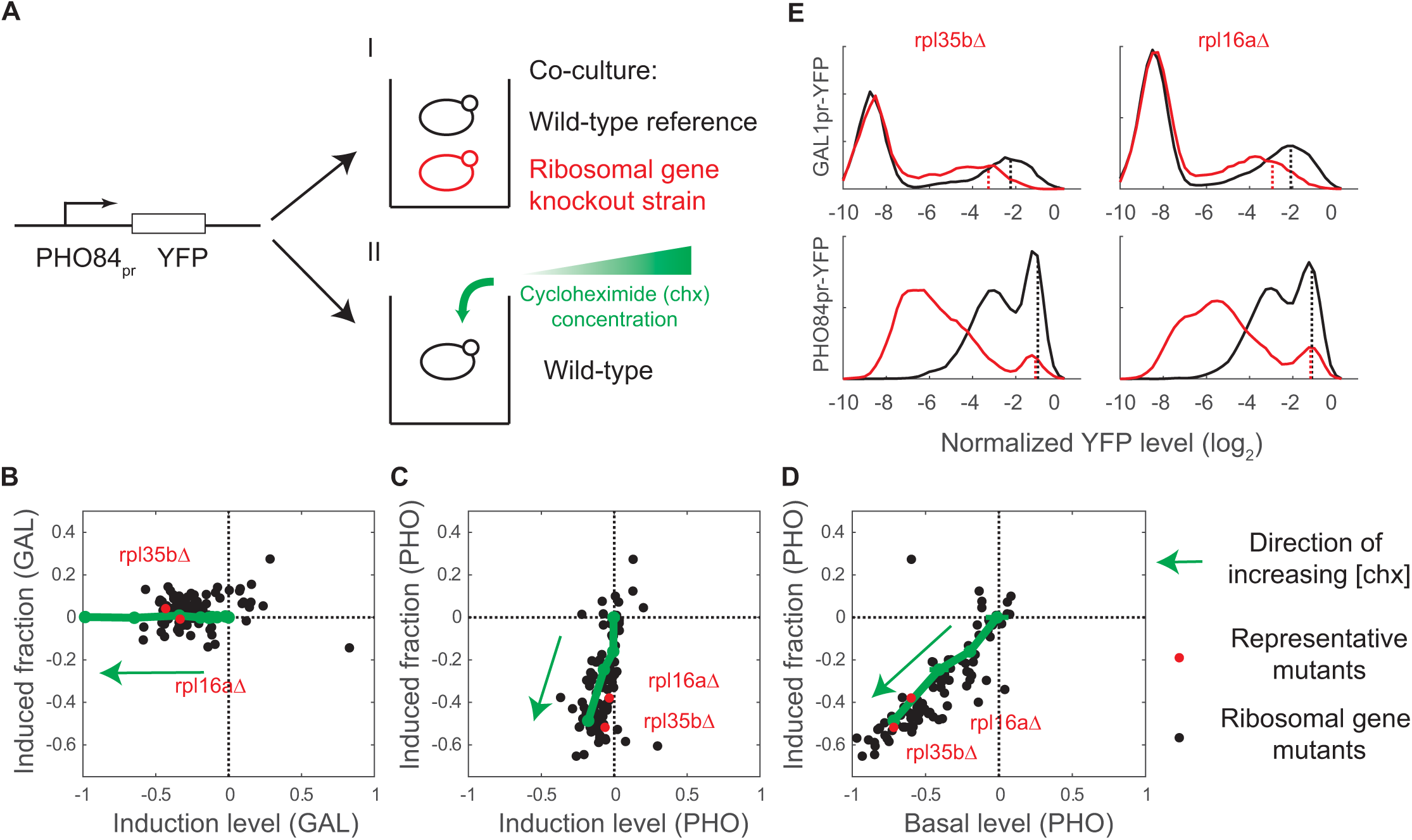
Effects of protein synthesis perturbation on the phosphate response (PHO) are distinct from the effects on the galactose response (GAL) (**A**) Schematic of experiment to quantify the effects of perturbing protein synthesis on the PHO response. A PHO84_pr_-YFP reporter was used to quantify PHO pathway activation in single cells. Protein synthesis was perturbed by either (I) knocking out genes involved in protein synthesis or (II) treating our wild-type strain with a titration of cycloheximide. (**B-D**) The effects of perturbing protein synthesis are different between the GAL and PHO response. Perturbation phenotypes were quantified by: 1) induced fraction, 2) induction level and 3) for the PHO response, basal expression level. A set of 95 strains each deleted for a gene involved protein synthesis (black dots) was assayed (GAL in **B**; PHO in **C** and **D**). Cycloheximide (chx), a protein synthesis inhibitor, was added at 11 different concentrations to a wild-type strain (green dots; green arrow denotes direction of increasing chx concentration). Cycloheximide has a dose-dependent affect on both the GAL and PHO response that phenocopies the effect of protein synthesis mutants. (**E**) The expression distribution for two representative mutants, *rpl16aΔ* and *rpl35bΔ* in (red dots in **B-D**). The GAL1_pr_-YFP (top) and PHO84pr-YFP (bottom) distributions of *rpl16aΔ* and *rpl35bΔ* mutants are shown (red), together with the co-cultured wild-type strain (black). The induction level metric is denoted (dashed line). The induction level is not change in PHO (bottom) while it is in GAL (top).

### Extension to other sources of genetic variation through simulation

We next wished to determine to what extent our results generalize to genetic variation beyond the complete loss-of-function variants we experimentally measured. Genetic variation in natural population is more complex genetically than the deletion library we analyzed. To generalize our results to account for a broader range of genetic variation, we developed a model where we accounted for 1) other types of alleles, i.e. hypermorphs and neomorphs as originally proposed by Muller (Muller, 1932), 2) variable number of alleles per gene, and 3) variable allele frequencies in the population (Figure 6). While the actual molecular cause of the variation can come from many sources, e.g. single nucleotide polymorphisms (SNPs), copy number variation, and indels, for the purpose of understanding the genetic architecture, it is only important to understand the effect of the genetic change on the trait, and hence for simplicity we will refer to all genetic variants as SNPs. Additionally, we assumed all SNPs contribute linearly to the trait with no epistasis. This assumption is based on the fact that a linear model using all SNPs genotyped in human height GWAS can explain a large fraction of height heritability (Yang et al., 2015; 2010).

**Figure 6.**
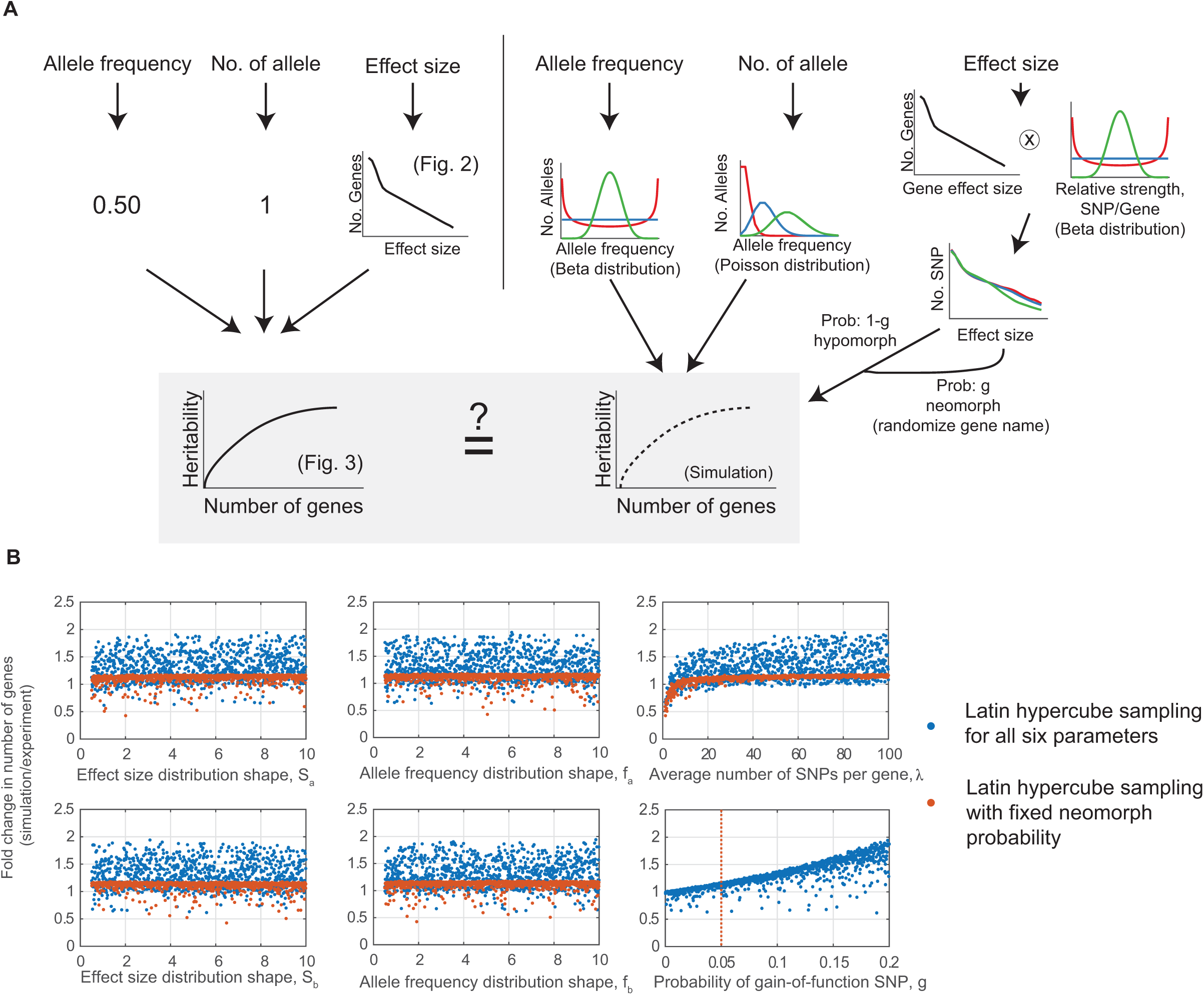
Effect size distribution estimated from gene deletions is informative for more complex genetic scenarios. (**A**) In figure 3, heritability versus gene number was estimated assuming an allele frequency of 0.5, exactly 1 SNP per gene, and the effect size distribution measured in figure 2. To simulate more complex biological scenarios, we sampled allele frequency from a Beta distribution, Beta(f_a_, f_b_); number of SNPs from a Poisson distribution, Poisson(λ); simulated hypomorphs by convolving the measured effect size distribution for amorphs with a Beta distribution, Beta(S_a_, S_b_); and neomorphs by randomly sampling from the hypomorph effect size distribution. The frequency of hypomorphs versus neomorphs was a constant, g, for each simulation. The extremes and middle of the range of each distribution are shown (red, blue, and green) (**B**) Comparison of the number of genes required to explain 80% of the heritability in the experimental and simulated data. Simulate data was generated by Latin hypercube sampling of the six parameters (1000 iterations; blue dots). The fraction of neomorphs (g) had the largest affect on the model. To examine the effect of the rest of parameters, g was set to 5% (vertical red dashed line), and Latin hypercube sampling was used was used to scan the remaining five parameter space 1000 times (red dots).

To instantiate the model (Figure 6) a series of functional forms and constants were assumed for each of the potential variables. The number of SNPs that affect a given gene was chosen from a Poisson distribution to reflect variable number of alleles observed in human genome (Sachidanandam et al., 2001). The effect size of hypomorphic SNP was modeled by multiplying a beta distributed random variable by the actual measured effect size of each affected gene. In this way, the maximum effect size was the complete loss-of-function and the minimum effect size was zero. A beta distribution was chosen to allow modeling of a wide range of different shaped distributions (Figure 6). To simulate neomorphic (gain-of-function) SNPs, we randomly selected a fraction SNPs, and reassigned their effect sizes with the effect size of randomly chosen SNPs. Lastly, the allele frequency for each SNP was chosen from a beta distribution.

In each simulation, we calculated the explained trait variation for each SNP, and then summed up all the SNPs for a single gene to obtain the explained variation for each gene. We then varied the parameters in each of the distributions of the variables introduced above. Specifically, we used Latin hypercube sampling to scan the parameter space of the distributions (blue dots, Figure 6), and then compared the number of genes that explain 80% of trait variation obtained from this model and our experimental results (Figure 3). The results from our simulation show variation in the fraction of neomorphs dominates variation in the model. However, as long as this fraction is below 5%, the results of the simulation do not vary from the experimental results by more than 17%. Neomorphic alleles are typically assumed to be rare. In order to determine the potential impact of the other parameters in our model, we fixed gain-of-function rate to be 5%. Resampling the other five parameters, we found that the average number of SNPs per gene is the second largest source of variation in our model. However, as long as on average 5 SNPs exist per gene in the population, the effect is negligible (orange samples in Figure 6B). Large-scale sequencing efforts have now identified ~20 million genetic variants in humans (Sherry et al., 2001). Even if 99% of these variants were neutral, there would still be enough SNPs per gene on average to support our conclusions. From this, we determined that our estimate of number of genes that influence a trait from our knockout data is largely insensitive to the parameters of our model, and quantitative analysis of complete loss-of-function alleles should be informative even for the analysis of less severe and of rare alleles.

## Discussion

In this work we sought to determine all the genes that can influence a pathway underlying a quantitative trait. Depending on the number of genes and the magnitude of the effect, pathways could in principle have a centralized or distributed architecture (Figure 1). To address this question we determined the effect size distribution of deletion mutants for four quantitative traits in yeast. We did this by measuring the response to galactose at the single-cell level for each deletion strain from the yeast library and by reanalyzing two additional quantitative screens that measure competitive growth rates (Breslow et al., 2008) and the unfolded protein response (UPR) (Jonikas et al., 2009) in the deletion library. We found that in all four cases, the distribution of effect sizes is such where a quarter to half of the genes follow an exponential distribution, with the rest of the genes having a negligible effect size (Figure 2). Based on a simple model to calculate heritability, we found this result implies that a large number of genes (5-9% of all genes) would be needed to cumulatively explain at least 80% of trait variation (Figure 3). Our results imply that there is a significantly larger subset of genes that affect each trait than previously appreciated, but that individually their effect is difficult to detect by less quantitative experimental methods. The results provide evidence for pathways having a distributed, rather than centralized, genetic architecture.

A distributed pathway architecture suggests that many genes that are not typically considered part of a pathway, such as the GAL pathway, could still play an important role in pathway function. We found that these “pathway modifiers” were enriched in several core cellular processes (Figure 4). Given the pleiotropic nature of these processes, it is not surprising that these genes can influence multiple traits. Unexpectedly, however, we found that a mutant in a core cellular process can have trait-specific consequences; e.g. a ribosomal mutant affects the induction level in the GAL response, but the induced fraction in the PHO response (Figure 5). This implies that, instead of making cells ‘sick’, biological processes underlying quantitative traits are likely affected by a large number of inputs that have the potential to act in a trait-specific manner.

Previous work had found that yeast traits were affected by fewer genes than we report here. Work by Bloom et al. used linkage analysis to identify quantitative loci underlying 46 yeast traits, and found a median of 12 loci affected each trait (Bloom et al., 2013). While it is possible that our four traits happen to be more complex than the traits that were analyzed by Bloom et al., we believe the differences result from the applied methods. If either the two yeast strains used in the linkage analysis of Bloom et al. are more related than two random isolates in a natural population or if the traits analyzed were under strong selection, this would lead to an underestimation of the number of genes. Because we are using a deletion library, we avoid the confounding effect of selection and the biases due to the limited number of alleles between two natural isolates. We therefore believe that the discrepancy between the results of these two works is at least in part due to the applied methods, in particular selection on growth rates.

### Mendelian vs. quantitative trait

A distributed genetic architecture, as observed in this study, has implication for patterns of genetic inheritance. Different individuals can have different numbers of alleles and the effect size of the strongest alleles can be different. Therefore, the expectation should be that the same trait, when examined in a pairwise manner between many individuals, should exhibit a range of segregation patterns from Mendelian to quantitative depending on the number and strength of the alleles. Indeed, this exactly what was recently observed for multiple traits in crosses between yeast strains (Hou et al., 2016). Furthermore, one should expect a smaller number of genes that contribute to a quantitative trait will have rare alleles that make the trait behave as a Mendelian trait. Indeed, this has also been found that many quantitative loci associated with normal human height variation contain genes underlying syndromes characterized by abnormal skeletal growth (Lango Allen et al., 2010).

### Application to human genetics

Our results suggest that the number of genes that can influence a trait, when extrapolated to humans, is ~ 1500. However, our results were focused on a single-celled microbe that has a more compact genome with a smaller number of protein-coding genes than metazoan genomes. To what extent might our observations generalize to human genetic variation, given the differences in genome architecture and complexity? Do human traits, especially ones involved in important human disease, also have such a distributed underlying genetic architecture? One way to assess whether the genetic architecture of yeast and human traits is different is to compare the number of genes and their corresponding effect size distribution.

While a small number of human diseases or traits can be explained by a small number of causative genes, e.g. three genes explain 50% of the genetic risk in macular degeneration (Maller et al., 2006), many traits are poorly explained by a small number of genes. For example, a GWAS on human height found that 423 loci explained less than 20% of total heritability (Wood et al., 2014). Similarly, 163 loci only explain 14% of heritability in Crohn’s disease (Jostins et al., 2012), and 100 loci, excluding major histocompatibility complex, explain less than 6% of heritability in rheumatoid arthritis (Okada et al., 2014). Since the explained fraction of heritability is far less than 100% in all these studies, it is difficult to accurately estimate the number of loci required to explain a majority of heritability in a human trait, but a reasonable estimate would be in the thousands. This suggests that the fraction of genes involved in a quantitative trait is similar in yeast and humans.

While the effect size distribution of human traits is poorly defined it is consistent with our results. Park et al. devised a method to determine the effect size distribution by taking into account all identified alleles and the power to have detected these alleles. From this they concluded that the effect size distribution alleles affecting human traits are monotonically increasing (Park et al., 2010). The range of possible distribution discussed in that work is consistent with an exponential distribution. While there is no good human data exists on the distribution of small effect size alleles, gene essentiality can be used as a rough comparison of the relative distribution of strong effect size allele between yeast and humans. Further supporting the similarity in effect size distributions, the number of essential genes in yeast and humans is similar. In total we believe this supports the idea that while the human genome is more complex than yeast, differences in genetic architecture are likely subtle and quantitative not large and qualitative.

### Implication of a distributed genetic architecture on human disease

High-throughput genetic interaction maps have suggested that cellular processes are deeply interconnected (Costanzo et al., 2016; 2010). But, it was not determined whether these connections were strong enough to be physiologically relevant. Our results demonstrate that cumulatively many genes of small effect size can make significant contributions to quantitative traits. Importantly, the effect sizes of these variants are not infinitesimal, and therefore we believe that increased power in GWAS would likely capture a significant portion of the missing heritability. This conclusion is consistent with work from Yang et al., which has shown that human genetic variants tagged in GWAS on body mass index is capturing the vast majority of heritability even if it is underpowered to identify the causative loci (Yang et al., 2015). Of course, increased power alone will not help identify which SNPs within a locus is causative.

Given that so many genes can affect a trait, a second expectation is that causative small effect size loci should be shared between many but not all traits. Indeed, correlation among genetic variants has been observed in a recent study using 24 human traits (Bulik-Sullivan et al., 2015). Interpreting these results has been challenging as these genetic correlations could arise from either a direct causative link between the two diseases or shared genetic factors. Our results suggest that these correlations can result from shared genetic factors that are enriched in core cellular processes. This means that there could be power in searching for processes that are significantly enriched between diseases that wouldn't typically be thought of as related. Finally, the spectrum of defects seen in some complex diseases could arise from the specific combination of small effect alleles in each individual.

In summary, our work provides a system-level perspective into the architecture of a quantitative trait. In contrast to most other works that focused on existing genetic variants, our work quantitatively determined the contribution of loss-of-function alleles. With further development of gene editing technologies and disease models, it will be interesting to test these conclusions in more complex systems.

## Author contribution

B.H. and M.S. designed the experiments. B.H. performed the experiments. B.H. and M.S. analyzed results and wrote the manuscript.

## Acknowledgments

The authors thank Rebecca Ward, Yarden Katz for critically reading the manuscript; the Springer lab for helpful discussions; and Shervin Javadi and Stratedigm for flow cytometry assistance. B.H. and M.S. are funded by National Science Foundation Grant 1349248; and by National Institutes of Health Grant RO1 GM120122-01. The authors have declared that no competing interests exist.

## Methods

### Re-analysis of quantitative screening that used the yeast deletion collection

Genome-wide screens that used the yeast deletion collection (reviewed in Giaever 2014) were re-analyzed. After downloading available effect size measurements for individual mutants, the measurement error of each assay in Table S4 was determined as the standard deviation of the differences of replicate measurements for identical strains (see Supplemental Information for details). The effect sizes were compared to measurement noise distribution, ~***N*** (0, measurement noise), to assign p-values for mutants. False discovery rates (FDR) were used to correct for the multiple hypothesis test problem. Significant mutants were defined as ones with FDR less than 0.5%.

### Plasmid and strain construction

We constructed a plasmid containing the GAL1 promoter driving YFP with a Zeocin resistance marker all flanked by regions that are homologous to the HO locus (A65V). This plasmid was digested with Not1 and transformed into the parental SGA strain (B56Y, MATx ura3Δ leu2Δ his3Δ met15Δ can1Δ::ste2pr-spHIS5 lyp1Δ::Ste3pr-LEU2 LYS2+cyh2)(Tong and Boone, 2006) to construct a base strain (D62Y), which was used to create both query and reference strains used in the GAL screen. Query strains (library SLL14) were constructed using the SGA techniques(Tong and Boone, 2006) on the deletion collection and base strain D62Y. A reference strain (F59Y) was constructed by a second transformation with a TDH3pr-mCherry construct. The PHO84 promoter driving YFP reporter (E40B) was constructed using similar method by using PHO84pr PCR-ed from FY4 (using primer CGTACGCTGCAGGTCGACGGATCCCGTTTTTTTACCGTTTAGTAGACAG and TAATTCTTCACCTTTAGACATTTTGTTATTAATTAATTGGATTGTATTCGTGGAGTTTTG) instead of the GAL1 promoter. The resulting PHO library (SLL15) and reference strain (I32Y) were used in the PHO screen.

### Galactose induction assay

Mutant strains from the deletion library that contains GAL1pr-YFP reporter and the corresponding reference strain were pinned onto YEPD agar plate before being inoculated into synthetic complete 2% raffinose medium to allow growth till saturation. Mutants and the reference strain were pinned together into 150 μl of fresh raffinose medium and grown for another seven hours, before being inoculated into 150 ul of synthetic complete 0.2% glucose and 0.3% galactose. After induction for eight hours, 10 ul of cultures were analyzed by flow cytometry LSRII with HTS. Each plate ran for ~20 minutes on the instrument. To ensure that all mutants underwent roughly the same induction time, no more than four plates were inoculated at a time. The induction level and induced fraction trait were based on measurements from two biological replicates in two separate days. Data were analyzed using a Matlab script (for representative raw data, see Figure S6).

### Phosphate starvation induction assay

Mutant strains from the library that contain the PHO84pr-YFP reporter and the corresponding reference strain were pinned on YEPD plate before being inoculated into synthetic glucose medium (SD). Mutants and the reference strain were then co-cultured in SD for 12 hours before washing in water twice and transferred into induction medium - synthetic glucose medium supplemented with 200 μM of K_2_HPO_4_. Medium recipe is from Wykoff et al. (Wykoff et al., 2007). Cultures were analyzed by a Stratedigm S1000EX cytometer cytometry. The three PHO traits were based on measurements from two biological replicates in two separate days. Data was analyzed using Matlab scripts.

### Fitting the effect size distribution

As the measured effects of most strains are close to measurement error, we first analyzed the effect size distribution of strains with significant measured effect sizes (FDR<0.5%). Mutant effect sizes were binned and fitted to exponential distributions. The only fitting parameter is the scale of the exponential distribution, which was estimated by maximizing the following log-likelihood function.

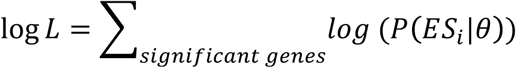
where *ES_i_* is the effect size of the *i^th^* significant gene, and *θ* is the the scale of the exponential distribution. The probability distribution is an exponential distribution defined over a range of effect size, i.e.:

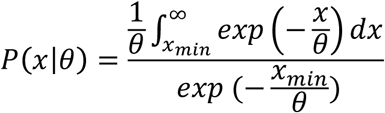

Parameters that maximize the likelihood of measurements were used for Figure 2. The fitted exponential distribution was extrapolated into the small effect size region to estimate the number of genes that are likely to follow the distribution. As a parsimonious model to explain our effect size measurements for four traits, we assumed the rest of mutants to have effect sizes as zero. Using this model, we predicted the expected effect size measurement distribution by convolving the true effect size distribution with the measurement noise of each assay (solid blue line in Figure 2). This distribution was then randomly sampled in 10,000 simulations and the standard deviation of the simulation was used as the confident zone of our estimation.

### Extrapolate the number of genes that affect quantitative traits to human traits

The number of significant genes were corrected by a factor determined by the gene number ratio between known human genes and screened yeast genes. The number of human genes is estimated as 22,500 (Pertea and Salzberg, 2010). The number of screened yeast genes was determined as the number of genes that passed quality control.

### Simulation of potential biases from the study of amorphs

In our model, we defined the explained heritability as the total explained heritability by all SNPs that affect each gene. As described in the main text, we simulated the number of SNPs that affect each gene as a Poisson distribution. The allele frequency and relative effect size are modeled using beta distribution. Gain-of-function SNPs were modeled by re-assigning effect sizes of a fraction of all SNPs by randomly sampling from the effect size distribution of all SNPs. In our Latin hypercube sampling, parameters in the two beta distributions ranged from 0.5 to 9, the fraction of gain-of-function SNPs ranged from 0 to 50%, and the average number of SNP per gene ranged from 1 to 100.

The heritability of each SNP is modeled as 2*S^2^*f*(1-f), where S is effect size and f is allele frequency. Measured knockout effect size on induced level is used in the model as complete loss-of-function effects. The code used for this simulation is available at Dryad.

### Gene Ontology analysis

Genes that are significant for all four traits (FDR<0.5%) were used as a hit list; all the genes that passed quality control were used as a background list. Gene Ontology analyses were done using GO TermFinder (Boyle et al., 2004).

### Spatial clustering algorithm

Each gene was represented a 4-dimensional effect size vector using the effect size measured for each of the four yeast traits. Since different traits have different units, we normalized each dimension of the effect size vectors by its scale, which is defined as the root mean square of the effect sizes of all the genes that significantly affect that trait. For any gene set, we determined the similarity of their effects on four traits by 1) filtering out all genes that are not significant to any of our traits; 2) calculate the circular mean of the normalized effect size vectors (e) as: 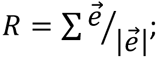3) calculate the circular deviation as *Var* = 1 – *R*. To determine the significance of this, we repeated the calculation 10,000 times after randomizing the gene names. Gene Ontologies that have at least five genes significant for any of the four traits were analyzed using the method above. Significantly clustered processes were defined as FDR < 0.01.

### Cycloheximide effect on GAL and PHO

Cycloheximide was purchased from Sigma (C7698). Cycloheximide was added directly to the induction media and this was the only change in the protocol from strains that were not exposed to cycloheximide. Cells were grown in a two-fold dilution series of cycloheximide with the highest concentration of cycloheximide being 20 μg/ml. Cycloheximide effects in Figure 5 were based on at least three biological replicates.

### Data and code Availability

All codes and raw data will be made available on Dryad.

## Supplemental figures and legends

**Figure S1.**
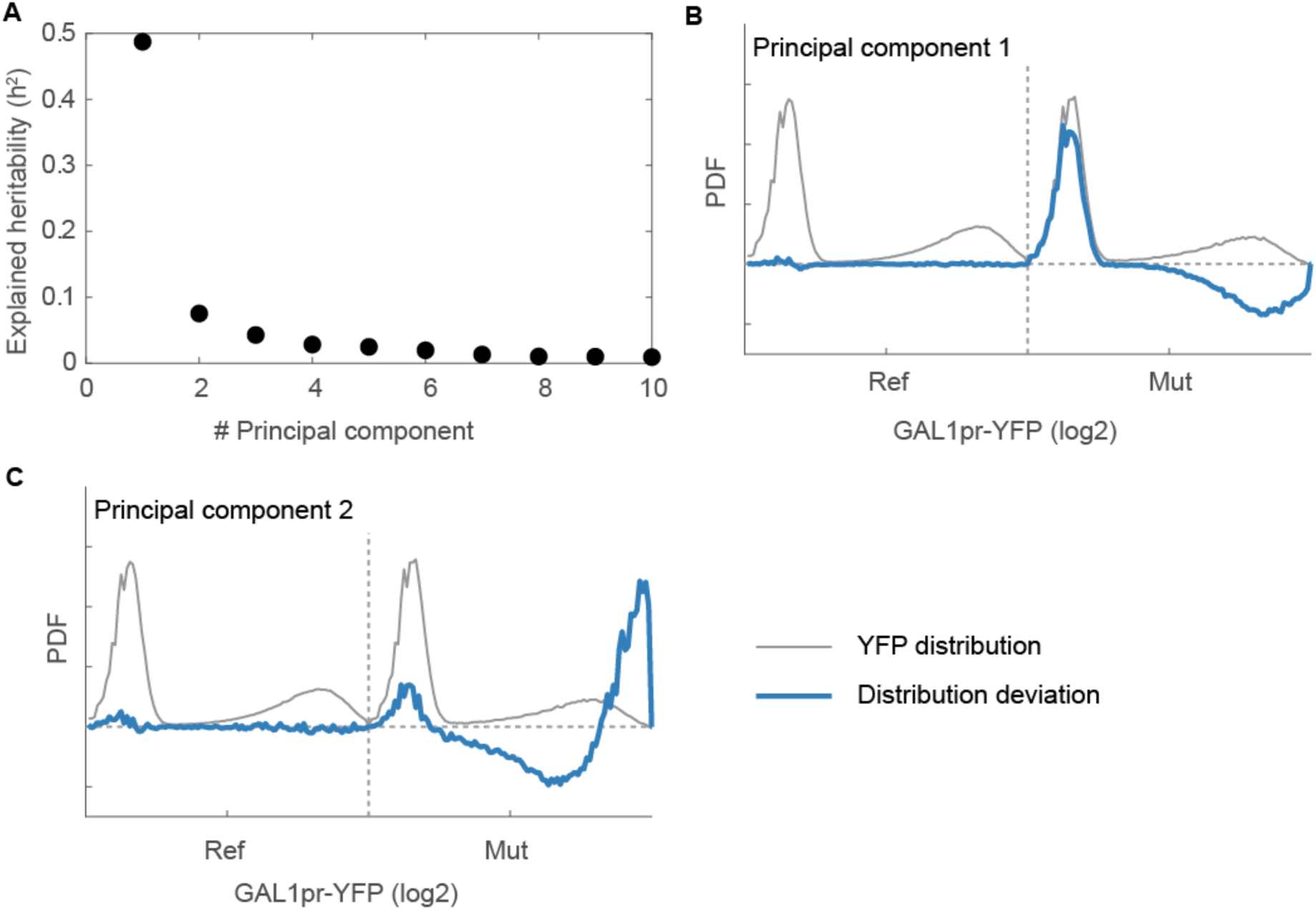
Determining modes of response with principal component analysis (Figure S1. Related to Figure 2) After data segmentation, histograms of GAL1pr-YFP for the mutant and reference strain for each sample were normalized, concatenated, and then analyzed using principal component analysis. (**A**) The fraction of variation explained by the first ten principal components. (**B-C**) Effects on GAL1pr-YFP distribution by the top two principal components. The average GAL1pr-YFP distribution of all reference and mutant strains are concatenated (gray). The principal component (blue) from the PCA analysis is the deviation from this average profile due to mutant effects. The horizontal line y=0 means no effects; i.e. the behavior of the wild-type strain. Note that the first two principal components correspond to biological properties, i.e. the induced fraction and induction level.

**Figure S2.**
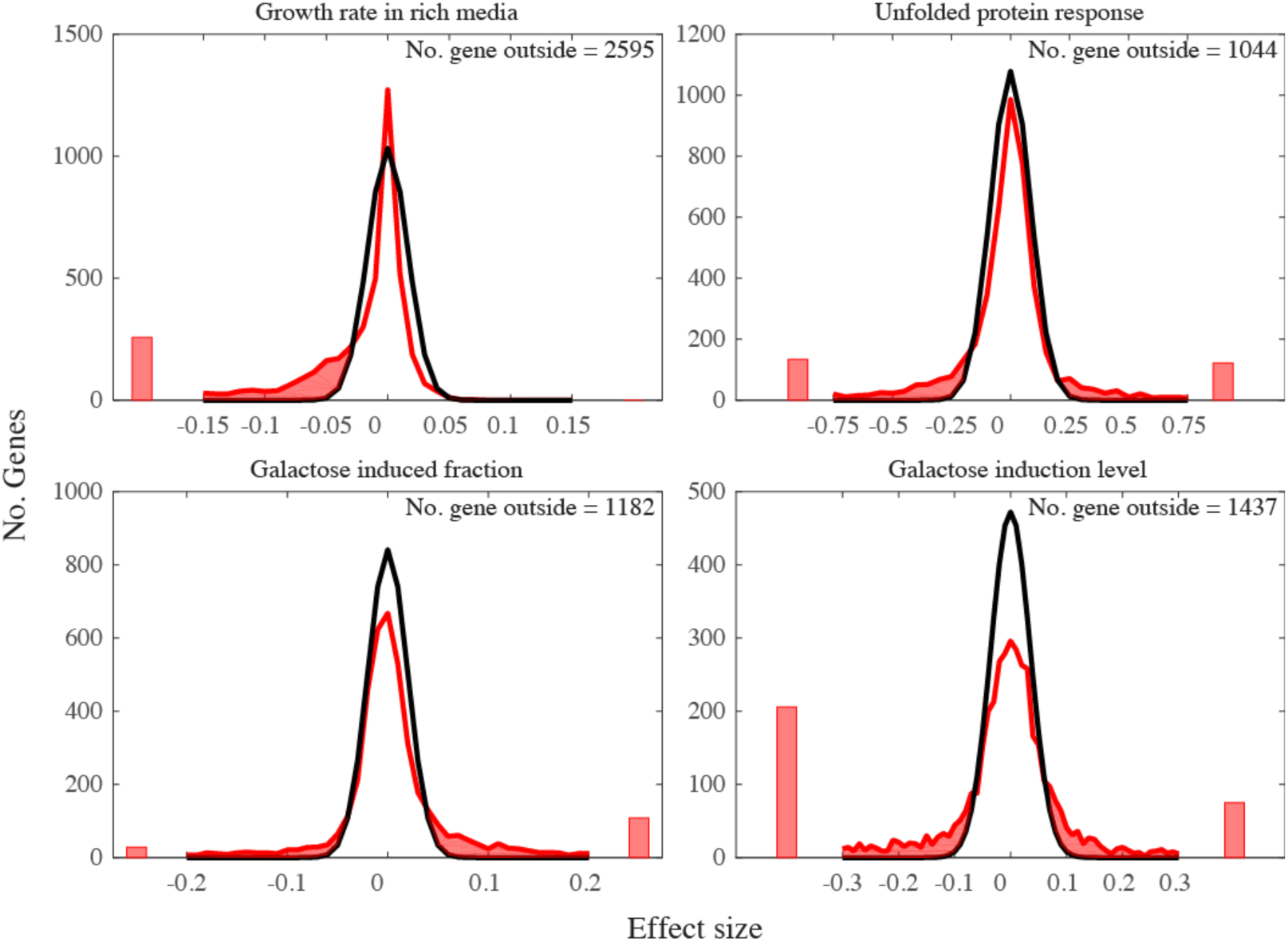
Effect size distribution versus measurement noise for four traits (Figure S2. Related to Figure 2) As many mutants have effect sizes that are close to or smaller than average measurement noise, the total number of genes that affects each quantitative trait was estimated by comparing the measured effect size distribution (red) and measurement noise effect size distribution (black). The measurement noise effect size distribution is the distribution of measurement noises between all replicated samples. The total number of genes that affect each trait was estimated from the number of genes in the shaded region.

**Figure S3.**
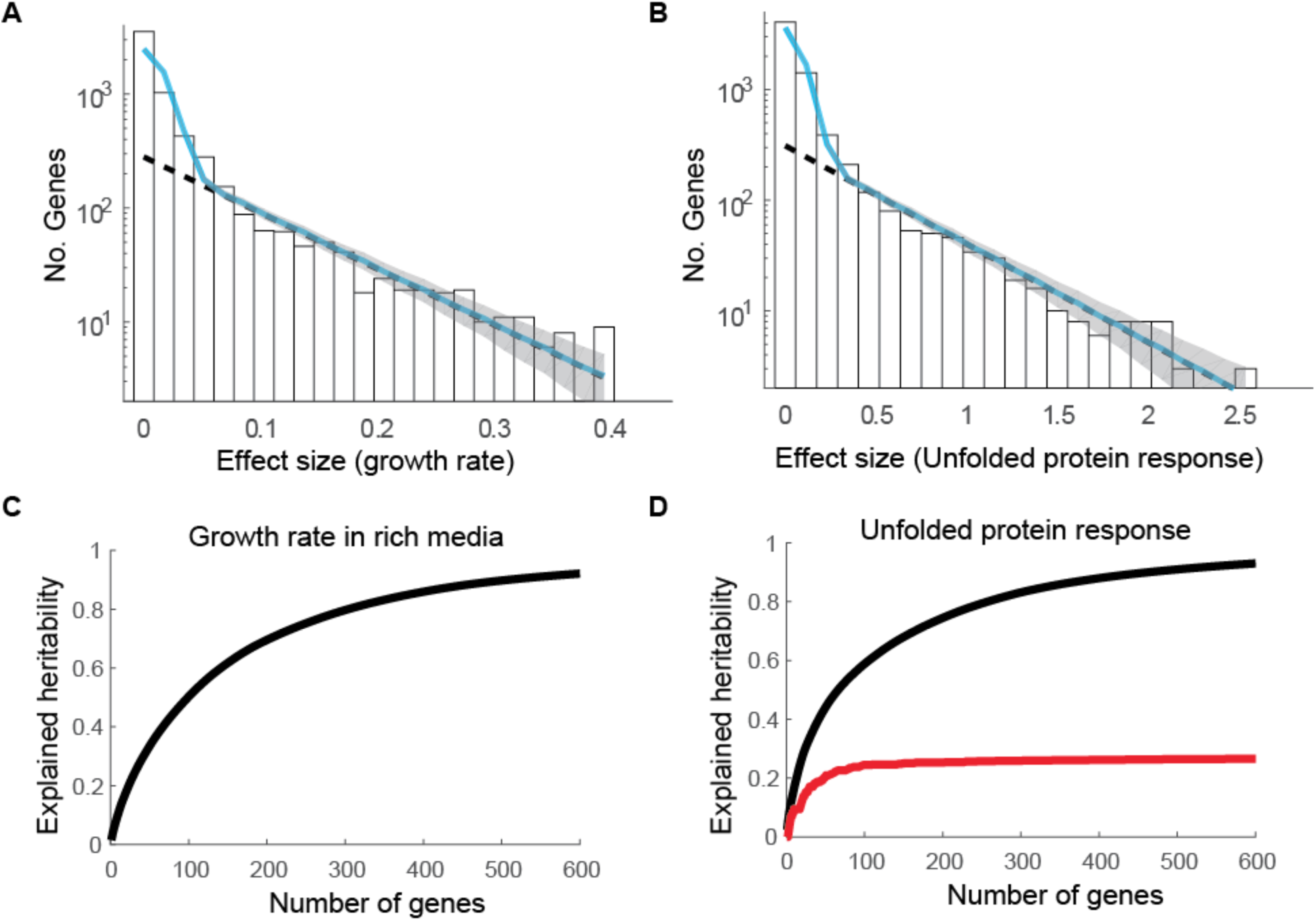
Reanalysis of two screens confirms that a large number of genes quantitatively affect yeast galactose response (Figure S3. Related to Figure 2; Figure 3) Data from two deletion studies (Breslow et al., 2008; Jonikas et al., 2009), one on growth rate in rich medium and one on the unfolded protein response (UPR), were reanalyzed. Both the effect size distribution (**A-B**) and explained heritability (**C-D**) were calculated as in **Figure 2** and **3**. Fit of the significant genes to an exponential (dashed line) has an R^^^2 of 0.91 for growth rate (**A**) and 0.94 for UPR (**B**). The fit of the full data to an exponential plus noise had an R^^^2 of 9.2 (**A**) and 0.95 (**B**). (**C-D**) The contribution to explained heritability, as calculated in **3A**, from UPR genes (red) or all genes (black) for growth rate in rich media (**C**) and UPR (**D**).

**Figure S4.**
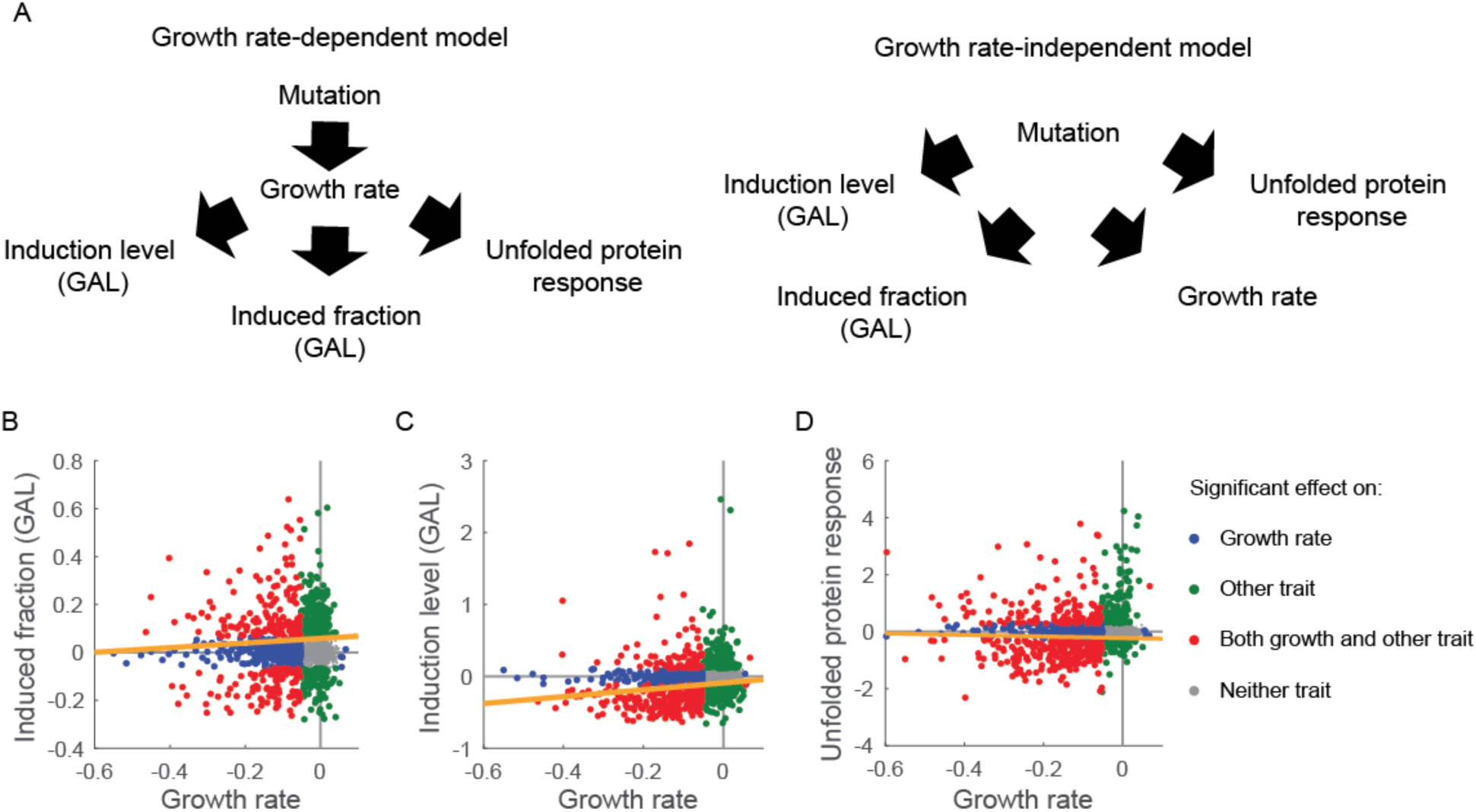
Affecting growth rate is not the sole mechanism for significant mutants to affect yeast GAL response and unfolded protein response (Figure S4. Related to Figure 4) (**A**) Two alternative models of how quantitative traits can be affected by gene deletion. In the growth rate-dependent model (left), mutants affect growth rate that in turn affects other traits. In the growth rate-independent model (right), mutants directly affect quantitative traits including growth rate. These two models can be distinguished by determining whether mutant effects on growth rate and other traits are correlated. (**B-D**) Mutant phenotypes for the unfolded protein response (**B**), GAL induced fraction (**C**) and GAL induction level (**D**) are plotted against the growth rate data reported by Breslow et al. Mutants were segmented into four quadrants based on whether the mutant had a significant effect (based on 0.5% FDR cut-off) on growth rate and non-growth rate trait: growth rate (blue), other non-growth rate trait (green), both (red), neither (gray). A linear fit of the points that are significant for both traits (red) is plotted (orange line).

**Figure S5.**
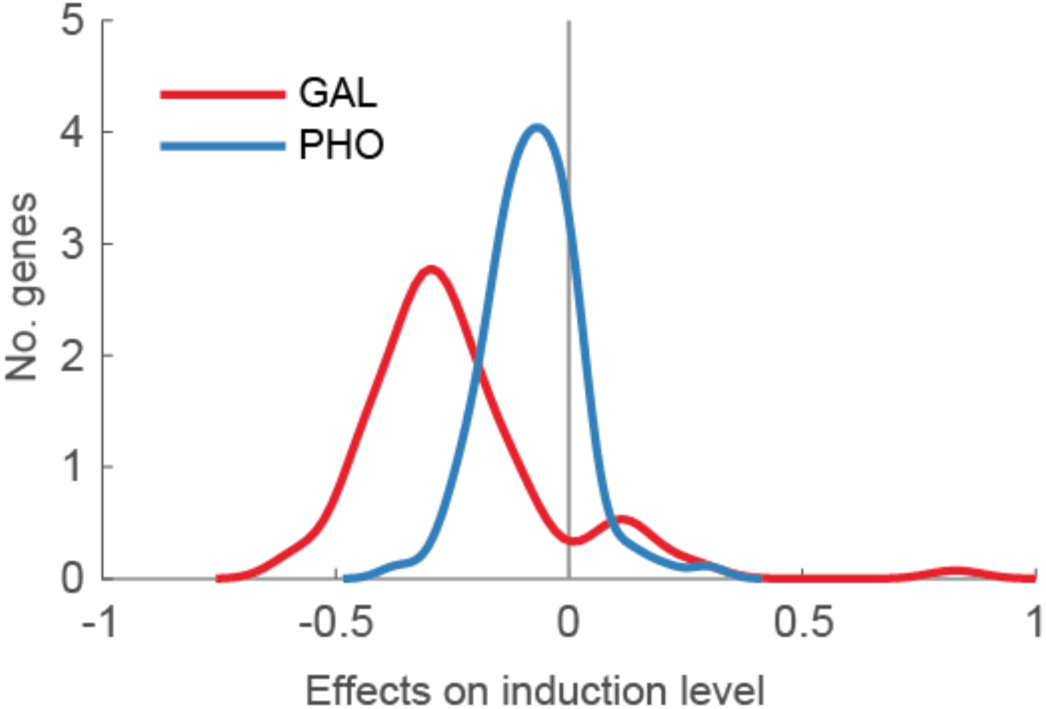
The difference of effects on GAL and PHO response by deleting genes involved in protein synthesis (Figure S5. Related to Figure 5) For each of the 95 mutants we tested that are involved in protein synthesis, the mutant effects on the induction level were quantified for the PHO (blue) and GAL (red) responses. The effect size distribution was smoothened with kernel smoothing with a bandwidth of 0.05. The two distributions are extremely unlike to have results from noise in a single distribution (p-value 3*10^-10^, two-tailed t-test). The magnitude of the average difference in effect size between the two distributions is 3 fold.

**Figure S6.**
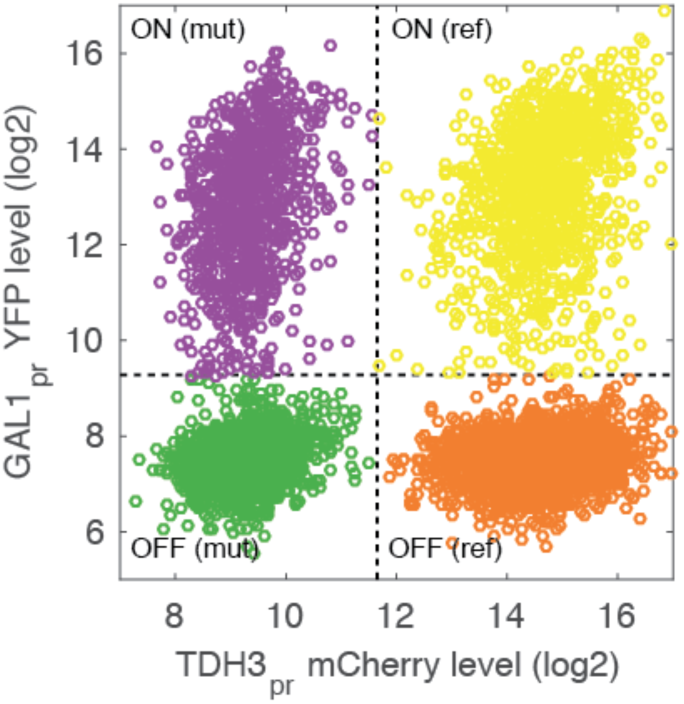
Data segmentation example (Figure S6. Related to Figure 2) An example is shown here using data from the first replicate sample for mutant *yal068cΔ*. Cell debris is filtered from the raw data using a FSC/SSC gate, and the mCherry vs. YFP values of remaining events are plotted. Data is segmented on the mCherry channel to separate reference and mutant strain, and on the YFP channel to separate the induced cells and uninduced cells. The horizontal and vertical dashed lines show the threshold used for segmentation.

## Supplemental tables and legends

**Table S1.** Enriched Gene Ontology for genes that significantly affect all four yeast traits (Table S1. Relates to Figure 4) GO TermFinder (Boyle et al., 2004) was used to analyze GO enrichment. The p-values are corrected for multiple hypotheses. Significant GOs are defined by the ones with corrected p-value less than 0.01.

| GO ID | GO Term | Corrected P-Value |
| --- | --- | --- |
| GO:0010467 | gene expression | 9.1E-21 |
| GO:0016070 | RNA metabolic process | 2.1E-15 |
| GO:0034641 | cellular nitrogen compound metabolic process | 2.8E-14 |
| GO:0006807 | nitrogen compound metabolic process | 2.4E-13 |
| GO:0090304 | nucleic acid metabolic process | 5.8E-13 |
| GO:0044260 | cellular macromolecule metabolic process | 1.6E-12 |
| GO:0044271 | cellular nitrogen compound biosynthetic process | 2.2E-12 |
| GO:0043170 | macromolecule metabolic process | 9.1E-12 |
| GO:0034645 | cellular macromolecule biosynthetic process | 1.2E-10 |
| GO:0009059 | macromolecule biosynthetic process | 1.8E-10 |
| GO:0006139 | nucleobase-containing compound metabolic process | 4.2E-10 |
| GO:0043933 | macromolecular complex subunit organization | 2.3E-09 |
| GO:0046483 | heterocycle metabolic process | 3.1E-09 |
| GO:0006725 | cellular aromatic compound metabolic process | 4.2E-09 |
| GO:1901360 | organic cyclic compound metabolic process | 1.4E-08 |
| GO:0044238 | primary metabolic process | 2.7E-08 |
| GO:0006396 | RNA processing | 1.2E-07 |
| GO:0044237 | cellular metabolic process | 1.2E-07 |
| GO:0044249 | cellular biosynthetic process | 3.1E-07 |
| GO:0002181 | cytoplasmic translation | 3.5E-07 |
| GO:0006355 | regulation of transcription, DNA-templated | 3.8E-07 |
| GO:1903506 | regulation of nucleic acid-templated transcription | 3.8E-07 |
| GO:2001141 | regulation of RNA biosynthetic process | 3.8E-07 |
| GO:0071824 | protein-DNA complex subunit organization | 4.1E-07 |
| GO:0010468 | regulation of gene expression | 5.8E-07 |
| GO:0044267 | cellular protein metabolic process | 6.7E-07 |
| GO:0051252 | regulation of RNA metabolic process | 7.2E-07 |
| GO:1901576 | organic substance biosynthetic process | 8.1E-07 |
| GO:0006351 | transcription, DNA-templated | 8.8E-07 |
| GO:0032774 | RNA biosynthetic process | 8.8E-07 |
| GO:0097659 | nucleic acid-templated transcription | 8.8E-07 |
| GO:2000112 | regulation of cellular macromolecule biosynthetic process | 1.2E-06 |
| GO:0071704 | organic substance metabolic process | 1.3E-06 |
| GO:0010556 | regulation of macromolecule biosynthetic process | 1.7E-06 |
| GO:0019219 | regulation of nucleobase-containing compound metabolic process | 2.0E-06 |
| GO:0009058 | biosynthetic process | 2.0E-06 |
| GO:0019538 | protein metabolic process | 2.1E-06 |
| GO:0031326 | regulation of cellular biosynthetic process | 4.3E-06 |
| GO:0009889 | regulation of biosynthetic process | 5.1E-06 |
| GO:0022613 | ribonucleoprotein complex biogenesis | 6.2E-06 |
| GO:0034654 | nucleobase-containing compound biosynthetic process | 7.4E-06 |
| GO:0051171 | regulation of nitrogen compound metabolic process | 8.4E-06 |
| GO:0006325 | chromatin organization | 9.1E-06 |
| GO:0008152 | metabolic process | 1.3E-05 |
| GO:0006412 | translation | 1.3E-05 |
| GO:0060255 | regulation of macromolecule metabolic process | 1.8E-05 |
| GO:0043043 | peptide biosynthetic process | 1.8E-05 |
| GO:0018130 | heterocycle biosynthetic process | 2.8E-05 |
| GO:0034728 | nucleosome organization | 3.3E-05 |
| GO:0043604 | amide biosynthetic process | 3.9E-05 |
| GO:0034660 | ncRNA metabolic process | 3.9E-05 |
| GO:0019438 | aromatic compound biosynthetic process | 4.0E-05 |
| GO:0019222 | regulation of metabolic process | 6.8E-05 |
| GO:0034622 | cellular macromolecular complex assembly | 6.9E-05 |
| GO:0080090 | regulation of primary metabolic process | 8.8E-05 |
| GO:1901362 | organic cyclic compound biosynthetic process | 9.1E-05 |
| GO:0034470 | ncRNA processing | 9.3E-05 |
| GO:0016568 | chromatin modification | 1.0E-04 |
| GO:0042254 | ribosome biogenesis | 1.1E-04 |
| GO:0051276 | chromosome organization | 1.1E-04 |
| GO:0006364 | rRNA processing | 1.2E-04 |
| GO:0031323 | regulation of cellular metabolic process | 1.3E-04 |
| GO:0043486 | histone exchange | 1.3E-04 |
| GO:0006518 | peptide metabolic process | 1.4E-04 |
| GO:0071840 | cellular component organization or biogenesis | 2.4E-04 |
| GO:0030490 | maturation of SSU-rRNA | 3.5E-04 |
| GO:0016072 | rRNA metabolic process | 4.7E-04 |
| GO:0071822 | protein complex subunit organization | 4.8E-04 |
| GO:0043603 | cellular amide metabolic process | 7.3E-04 |
| GO:0065003 | macromolecular complex assembly | 1.1E-03 |
| GO:0044085 | cellular component biogenesis | 1.1E-03 |
| GO:0043044 | ATP-dependent chromatin remodeling | 1.3E-03 |
| GO:0010629 | negative regulation of gene expression | 1.7E-03 |
| GO:0045892 | negative regulation of transcription, DNA-templated | 2.0E-03 |
| GO:0051253 | negative regulation of RNA metabolic process | 2.0E-03 |
| GO:1902679 | negative regulation of RNA biosynthetic process | 2.0E-03 |
| GO:1903507 | negative regulation of nucleic acid-templated transcription | 2.0E-03 |
| GO:0000462 | maturation of SSU-rRNA from tricistronic rRNA transcript (SSU-rRNA, 5.8S rRNA, LSU-rRNA) | 2.1E-03 |
| GO:0006338 | chromatin remodeling | 3.7E-03 |
| GO:0042274 | ribosomal small subunit biogenesis | 3.8E-03 |
| GO:0010558 | negative regulation of macromolecule biosynthetic process | 4.0E-03 |
| GO:2000113 | negative regulation of cellular macromolecule biosynthetic process | 4.0E-03 |
| GO:0016569 | covalent chromatin modification | 5.0E-03 |
| GO:0016570 | histone modification | 5.0E-03 |
| GO:0006357 | regulation of transcription from RNA polymerase II promoter | 5.7E-03 |
| GO:0051172 | negative regulation of nitrogen compound metabolic process | 8.8E-03 |
| GO:0031327 | negative regulation of cellular biosynthetic process | 9.4E-03 |
| GO:0045934 | negative regulation of nucleobase-containing compound metabolic process | 9.6E-03 |

**Table S2.** The number of genes that affect growth rate and each of the three non-growth traits (Table S2. Relates to Figure 4) Genes that significantly affect the unfolded protein response, induced fraction (GAL), and induction level (GAL) were compared to the genes that significantly affect growth rate. The total number of genes that were measured in both growth rate and the other trait is listed. Of this total number, the number that significantly affected growth rate, significantly affected the non-growth rate trait, and significantly affected both traits is listed.

|  | No. genes also<br>assayed in growth<br>rate screen | No. genes<br>significantly affect<br>growth rates | No. genes<br>significantly affect<br>the non-growth rate<br>trait | No. genes<br>significantly affect<br>both |
| --- | --- | --- | --- | --- |
| Unfolded protein<br>response | 4152 | 779 | 594 | 369 |
| Induced fraction<br>(GAL) | 3869 | 634 | 595 | 254 |
| Induction level (GAL) | 3869 | 634 | 744 | 316 |

**Table S3.Significantly spatially clustered Gene Ontology (Table S3. Relates to Figure 4)**

For each Gene Ontology that is spatially clustered, the direction in the four-trait space is shown, as well as the p-value and false discovery rate (FDR). Significant GOs were defined by FDR < 0.01.

**Table S4.**
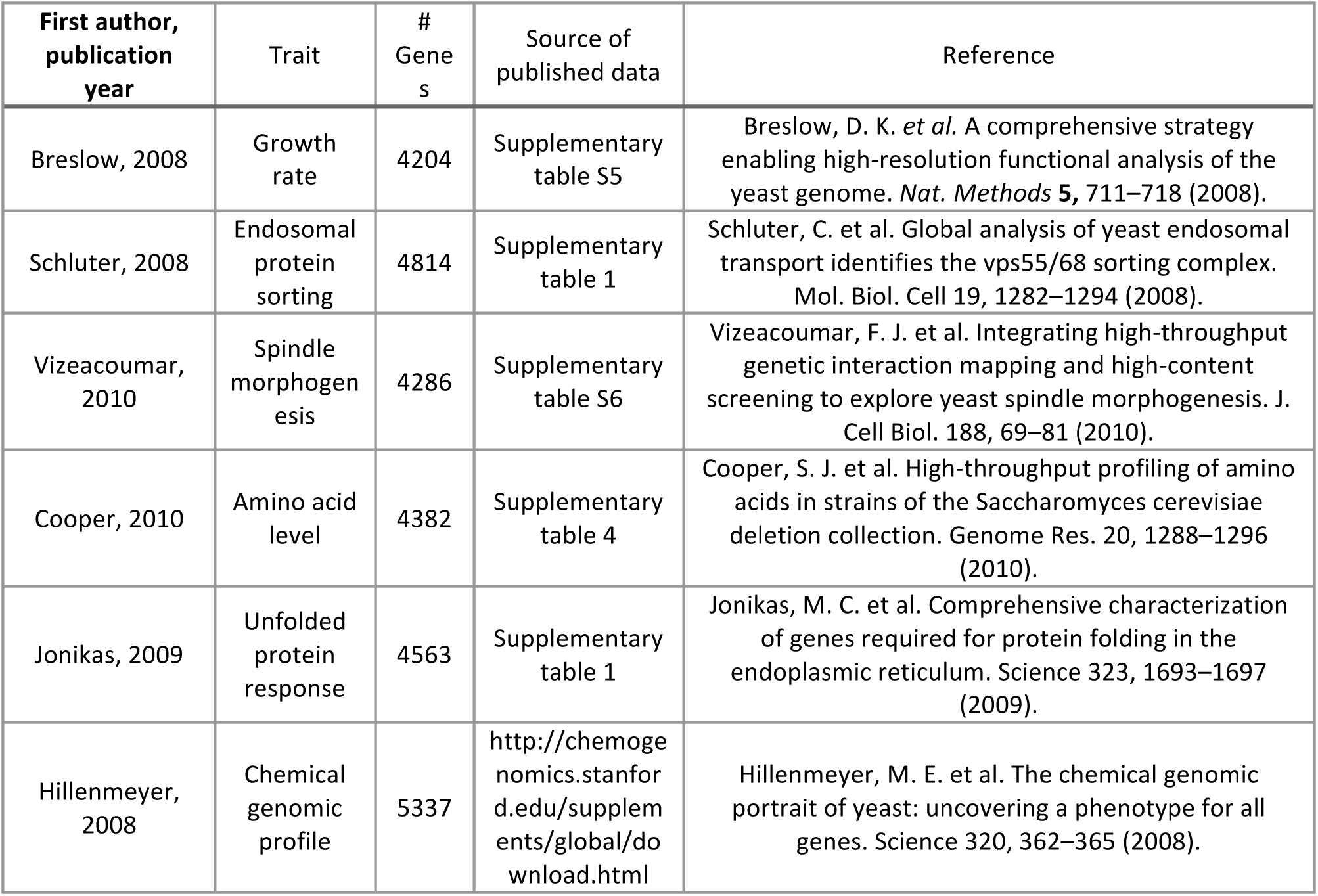
Quantitative screens that are analyzed for gene effect size distribution (Table S3. Relates to Figure 2) We manually scanned over 200 published deletion library screens to identify datasets that could be reanalyzed to potentially determine an effect size distribution. Of these 200 papers, we found only 6 that contained datasets in a form that was suitable for our reanalysis.

## Supplemental Text

### Re-analysis of previous quantitative screening using yeast deletion collections

Since the release of the yeast deletion collection, a large number of studies have been performed (Giaever and Nislow, 2014) potentially providing a rich source to understand the quantitative effects of gene deletions on traits. Unfortunately, the raw data was not published and readily available for all but a small handful of these studies (**Table S4**).

Data from each screen in the **Table S4** was analyzed using the following method: 1) download raw data; 2) determine the measurement error; 3) calculate p value for each gene by comparing effect size measurement to measurement error (two-tailed t-test, assuming measurement error is Gaussian distributed); 4) correct the p values for multiple hypothesis tests by calculating false discovery rate; 5) identify the number of significant genes as ones with FDR<0.5%.

The measurement error for individual assays were determined as below. We assume that the true effects of deleting the *i^th^* gene is *x^i^*. The two independent measurements, *x^i^*,*^j^* = *x^i^* + ^*ϵi*,*j*^ for *j* = 1,2, where *ε* is the measurement noise term. Assuming that measurement noise follows a Gaussian distribution, i.e. *ϵ*^*i*,*j~N*^(*0, σ*). The difference of the two measurements on the identical strain will reveal information about the standard deviation of measurement noise. Specifically, since

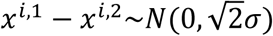
we can derive the following estimate of the standard deviation of measurement noise:

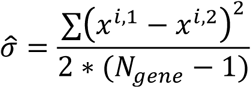
This method was applied to the raw data from the six assays in **Table S4**. Breslow et al. had different number of replicates (Breslow et al., 2008), and hence the measurement error for the individual mutants varied depending on the number of replicates. Specifically, among 4204 assayed genes, 2809 genes have one measurement, 874 genes have two replicates and 521 genes have at least three replicates. To avoid this complication, we only used the data from the first measurements and used the remaining data to estimate the measurement error. We first estimated the measurement error by applying the equation above to the replicate measurements of 874 strains with two measurements and determined measurement error as 0.015. Then we calculate the measurement error the 521 strains for which three measurements had been made. This yielded a measurement error of 0.017. As these two estimations are close, we use the average (0.016) as the measurement error for the assay. We observed that the measurement noise tends to be larger for strains with large effect size, which means that most strains with moderate effect sizes probably have smaller than estimated measurement error. Hence, we do not believe that this method will overestimate the number of genes affecting the growth rate trait.

Similarly, mutants in Jonikas et al. had different numbers of replicates (Jonikas et al., 2009). Measurement noise decreased as the number of replicate increased. As a conservative estimate of effect size measurements, we treated all measurements as if they had only two replicate data. To estimate the measurement error, we used the data from 541 strains with exactly two replicate data. In the original paper, the standard deviation of measurements for each strain was reported. Since there were only two measurements for these strains, the standard deviation equals the half of the difference between two measurements. Assuming that measurement noise of each replicate data followed *N* (0, *σ*), the expectation of the half of the difference of two independent measurements is *σ/√π*. When plotting the histogram of this data, we found that a number of measurement have exceptionally large measurement error, which artificially increased our estimation. After removing strains with measurement error larger than 0.5, the resulting measurement standard deviation has an average as 0.0678. Hence, we estimated the measurement noise as 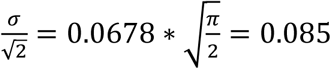.

Mutants in Schluter et al. were assayed in replicates for both haploid and diploid strains(Schluter et al., 2008). We applied the equation above to this data and determined that the measurement error was 0.027 for the MATa, haploids, 0.021 for the MATalpha haploids, and 0.042 for diploids. We used the average of these three to estimate measurement error (0.030). Vizeacoumar et al. provided p values for each mutants in the assay (Vizeacoumar et al., 2010). We convert the p value back to a z-score using Matlab function *norminv*(). While Copper et al. published raw data, they did so for only one replicate and hence we did not proceed with further analysis on this data set (Cooper et al., 2010).

Furthermore, we analyzed the raw data from Hillenmeyer by comparing the measured effect sizes in independent experiments using the same condition (drug name, dosage and the duration to apply the drug) in separate batches (Hillenmeyer et al., 2008). We found a large variation of the reproducibility between these replicates, determined as the pair-wise Pearson correlation coefficient (ranging from -0.2 to 0.99 with a median of 0.36 depending on the condition used). Hence we did not analyze the data further more.

To evaluate the number of gene deletions that significantly affected each of the quantitative trait, we first considered a null model where all gene deletions had no effects on the assayed traits. We expected the measured effect sizes to follow a normal distribution determined by measurement noise, i.e. *~N*(0, measurement noise). However, we found this was not the case for all the traits that we analyzed. To better illustrate this, we re-scaled the effect sizes by measurement error for each trait, and plotted the histograms of the re-scaled effect sizes for the gene deletions that have effect sizes at least 3-fold of the estimated measurement noise in **Figure S2**. We found the distributions were continuous. Note that only about (1-99.7%)*5000=15 genes were expected from the noise distribution. This suggests that the measured effects of most of the plotted genes were not from the measurement noise. Note that the data from Vizeacoumar was not shown here as the majority of genes have effects that are within three-fold of measurement noise.

To identify assays that are sensitive enough the measure the effect sizes of as many genes as possible. We estimated the number of genes that significantly affect each of the analyzed traits by comparing the measured effect sizes to measurement noise. Using a cutoff of FDR < 0.5%, we determined that two screens by Jonikas et al. and Breslow et al. are suitable for effect size distribution analysis as they have smallest measurement errors.

### Flow cytometry data processing

Raw data was exported from an LSRII or Stratedigm in fcs3.0 file format. All data was loaded using customized *MATLAB* code. In briefly, data from each sample was first filtered on FSC/SSC channel to remove cell debris, and on SSC channel to normalize for cell size. The FSC/SSC gates were drawn manually on pooled samples. The SSC gate was determined to include events between the 25^th^ to 75^th^ percentiles of the pooled sample. Pooled samples were also used to find thresholds on YFP and mCherry channels to segment induced vs. uninduced cells, and reference vs. mutant cells (**Figure S6** as an example). Mutants were filtered to ensure that there are at least 700 events for both reference and mutant cells in at least one biological replicates. Mutants in twelve plates in replicate one of the GAL screen have higher induced fraction than the reference strain in the same sample. Data from the second replicate were used for these mutants in the future analysis. For the PHO screen, we calculated the standard deviation of the effect size differences between two replicates for each of the three traits. The effect size measurements for fourteen mutants are greater than five-fold of these standard deviations. These strains were filtered from future analysis.

### Principal component analysis on reporter expression distribution of the entire deletion collection

Yeast responds to a mixture of glucose and galactose in a bimodal way. We measured expression level of GAL1pr-YFP in single cells for each of the mutant strain in the deletion collection. We generally observed that the reproducibility was higher when normalizing the distribution by comparing the mutant distribution to the reference distribution in the same well (see the section **Data Normalization** for details); as opposed to analyzing the mutant data directly. This is presumably due to slight variation between wells, plates, and days.

To find appropriate metric by which to analyze the mutant strains, we performed PCA analysis. We did this by pooling reference and mutant YFP distribution from two replicates. After data segmentation, the GAL1pr-YFP distributions of both reference strain and mutant strain were binned into 92 equally size log2 bins ranging from the maximum to minimum value. Data was normalized to probability distribution, separately for reference strain and mutant strain in each sample. PCA results were shown in **Figure S1**. The first three principle components explain ~ 60% variation. By manually examining the shape of each principle component, we could provide a plausible biological explanation for the major components. The first vector affects the induced fraction without affecting the expression level. The second vector has two effects, shifting the expression level of induced cells as well as changing the fraction of induced cells. The third vector change the expression level of both uninduced cells (basal level) and induced cells. In further analysis, we found that the expression level of uninduced cells could not be accurately determined for the majority of strains in our assay for GAL1pr-YFP reporter, and hence only the induced fraction and the induction level are used in the main text. This third metric was used for analysis of the PHO response.

### Data normalization

The induced fraction and induction level traits were calculated for each mutant strain using the following method. First, the induced fraction and induction level were calculated for reference strains and query strain in each sample. The induced fraction was calculated as the ratio of the number of induced events over the number of all events. The induction level was calculated as the average level of YFP of the induced cells. For both traits, the mutant value was regressed against the reference value using the *Matlab* function *robustfit*(). The residual of each measurement from the fit was averaged between two replicates to determine the final values of the induced fraction and the induction level.

### Estimate the number of genes that affect yeast quantitative traits

The noise distribution determined from measurement noise estimation was overlaid with the actual effect size measurements. Both curves were normalized to the total number of genes. The area of the region where the actual effect size distribution was outside the measurement noise distribution was determined for estimating the number of genes that affected each of the four yeast traits (**Figure S2**).

### Compare the number of detected mutants by using induced fraction and induction level vs. average expression level

Our screening data on the yeast galactose response provided a test for estimating the total number of significant mutants using different metrics. This is interesting as many biological traits could usually be defined in different ways, yet it was unclear to our knowledge how much potentially subtle differences in metric could influence genes identified. Here when we are referring to different metrics it is probably easiest to think of them as different sub measurements. For example, if one measured standing height as opposed to sitting height, would one uncover different sets of genes. In our case, the effect of gene deletion on galactose response can be represented as the two GAL traits as used in the main text, or alternatively we could simply use the average YFP level as used in Jonikas et al (Jonikas et al., 2009). To estimate such effects, we re-analyzed our data by quantifying not just the two GAL traits, but also the average YFP level. After applying the same method to detect mutants that significantly affect yeast GAL response, we found that the two-traits method detected more mutants (1104) than the average YFP method (593). In addition, the one-trait method could not reveal the distinct modes by which different mutants worked; i.e. 50% reduction in average can come because 50% of cells don’t induce or 100% of cells are 50% less induced. Hence our data suggested that, biological meaningful decomposition of a complex trait will increase detection sensitivity, and provides new biology insights to understand traits.

### Genes that saturated our assay

Our GAL assay was designed to detect genes of small effect size, and as a result, ten genes of larger effect size saturated our assay. These genes were manually verified by inspecting the YFP distribution of the raw data. These genes are: *GAL4* (*YPL248C*), *GCN4* (*YEL009C*), *GAL80* (*YML051W*), *GAL1* (*YBR020W*), *SNF3* (*YDL194W*), *STI1* (*YOR027W*), *REG1* (*YDR028C*), *GAL3* (*YDR009W*), *SNF2* (*YOR290C*), *HSC82* (*YMR186W*). This is important when calculating the explained heritability for top N genes (see main text). One of our main arguments is that the number of genes that affect a quantitative trait is around 8% of the genome. If the true effects of these ten genes is much larger than what we estimated, the number of genes that affect a quantitative trait could be smaller.

When using the nominal values of the measurements as effect sizes of these genes, we determined that the total contribution of these genes are 25.2% and 7.5% for induction level and induced fraction respectively. As another way to estimate the effect sizes of these genes, we randomly sampled the effect size distribution. The average contribution of these genes is 27.8% and 10.5% respectively, suggesting that this alternate method does not strongly affect conclusion.

### Overlapping among genes that are significant for each of the four studied traits

We examined the overlap between significant genes that affect growth rate and ones that affect each of the three other non-growth traits. To do so, genes with missing data in one of the data sets were removed. The result is in **Table S2**. The p-value was calculated between each pair of growth rate and non-growth rate trait, using a hypergeometric test (one-tailed).

### Compare the effects on GAL and PHO response by deleting genes involved in protein synthesis

For 95 genes involved in protein synthesis, we compared their effects on GAL and PHO traits in the main text and **Figure S5** using t-test (two-tailed). The average difference between the effects on GAL and PHO is 0.15. The standard deviations of effects on GAL and PHO are 0.21 and 0.10. As an alternative method to test for significance, we pooled the measured effects on GAL and PHO response and randomly split the pooled data into two groups for 1,000,000 times and calculated the difference between two groups. The observed difference (0.15) is not observed in the randomized sample. Hence we determined that p < 10^-6^ using this method.

### Canonical genes involved in galactose signaling and unfolded protein response

Glu/Gal gene list: *GPB2, IRA1, TOS1, GLK1, GPA2, GAL83, SAK1, GLC7, YCK1, BCY1, RGT1, ELM1, TPK3, HXK1, GPR1, RGT2, SNF3, REG1, MTH1, MSN5, SIP1, SNF1, MIG1, SNF4, SIP2, PDE1, HXK2, CYR1, TPK1, GRR1, SDC25, CDC25, SIP5, RAS2, YCK2, IRA2, STD1, RAS1, RGS2, PDE2, GPB1, TPK2, GAL1, GAL3, GAL80, GAL4, SNF2, GCN4, HSC82, STI1*

Gene localized in ER, Golgi, and early Golgi are (298 genes): *YEL031W, YJR117W, YFL025C, YJL062W, YML012W, YAL023C, YJR118C, YML055W, YML013W, YOR002W, YGL084C, YCR044C, YER122C, YNL219C, YNR030W, YDL095W, YML115C, YGL020C, YGL054C, YIL039W, YEL036C, YPL227C, YOL013C, YMR022W, YMR161W, YKL212W, YDL192W, YLR110C, YGL167C, YMR264W, YAL058W, YER083C, YDR027C, YLR372W, YCR094W, YLR268W, YNL238W, YMR307W, YJL029C, YBR171W, YDL100C, YGL226C-A, YBR106W, YJR073C, YNL322C, YGR229C, YGR284C, YJR010C-A, YML128C, YFR041C, YNL323W, YEL042W, YMR123W, YBR015C, YJR075W, YBR162W-A, YCR067C, YJL004C, YCR017C, YAL026C, YOR216C, YIL090W, YAL007C, YNL041C, YJL123C, YIL040W, YBR164C, YCL045C, YNL051W, YIR004W, YPL050C, YPL051W, YGL126W, YCR034W, YMR292W, YDR233C, YNL297C, YGL005C, YDR245W, YBR036C, YDR221W, YPL192C, YLL014W, YDR508C, YEL001C, YER005W, YDR137W, YDL099W, YGL231C, YHR108W, YMR238W, YAL053W, YIL027C, YER072W, YML038C, YER120W, YEL027W, YIL030C, YDR492W, YJR131W, YMR010W, YHR181W, YPR063C, YIL124W, YLR350W, YJR088C, YBL011W, YML048W, YNL044W, YDR358W, YOR311C, YDR411C, YMR272C, YNL049C, YMR015C, YDL052C, YJR134C, YKL096W, YNL280C, YLR194C, YER113C, YDR077W, YDR055W, YNR021W, YNL327W, YLR130C, YNR039C, YJL099W, YKL146W, YPR003C, YHL017W, YOR245C, YER166W, YBR132C, YOR016C, YPR090W, YNL300W, YLR250W, YGR038W, YPL259C, YPR071W, YKL065C, YKL046C, YPL274W, YEL048C, YOR317W, YDR100W, YNL146W, YMR253C, YJR031C, YER011W, YJL078C, YIL016W, YML037C, YGR247W, YFL004W, YBR023C, YIL044C, YMR052W, YDL204W, YBR067C, YDR153C, YIL043C, YNL095C, YDR476C, YOR307C, YOR321W, YCR011C, YMR237W, YMR071C, YER004W, YPR028W, YGL255W, YPL170W, YKL063C, YJL044C, YLR023C, YMR215W, YMR251W-A, YGR261C, YPR091C, YDR056C, YLL028W, YLR330W, YBL010C, YNR019W, YGL124C, YDR294C, YNL046W, YDR519W, YKR088C, YLR042C, YKL094W, YCR048W, YCR043C, YDR084C, YKR067W, YJL196C, YLL061W, YML101C, YDL232W, YOL030W, YMR054W, YDR410C, YBR273C, YLR120C, YHR110W, YOR044W, YDL137W, YJL171C, YOR285W, YMR029C, YLR064W, YPL137C, YOR092W, YBR159W, YGL083W, YNL156C, YDL128W, YBR296C, YOR175C, YJL198W, YOL101C, YHL019C, YJL117W, YGR263C, YML059C, YOR214C, YNR013C, YOR087W, YJL192C, YGR177C, YBL102W, YPL195W, YLL052C, YLR390W-A, YDR264C, YOR299W, YMR152W, YLL055W, YDR424C, YBR287W, YEL040W, YNL125C, YHL003C, YBR283C, YDL121C, YHR045W, YNR075W, YOR377W, YHR039C, YGL010W, YCL025C, YNR044W, YLR050C, YOL137W, YOL107W, YDL018C, YDR307W, YDR297W, YNL190W, YDR503C, YBR177C, YGR266W, YER019C-A, YLR034C, YOR322C, YGR260W, YDR349C, YJR015W, YPL246C, YMR058W, YBR290W, YLL023C, YDR205W, YHR123W, YJL024C, YJL212C, YLR292C, YPL207W, YKR027W, YIL076W, YBR288C, YJL183W, YKL008C, YJL207C, YML067C, YGR089W, YOR291W, YNL111C, YEL043W, YPL234C, YLR056W, YKL096W-A, YGR157W, YHR060W, YLR039C, YHR079C*

